# Recognition of distinct sleep states in *Drosophila* uncovers previously obscured homeostatic and circadian control of sleep

**DOI:** 10.1101/2025.06.23.661143

**Authors:** Lakshman Abhilash, Reed Evans, Orie Thomas Shafer

## Abstract

Understanding the mechanisms underlying homeostatic sleep regulation is a central unmet goal of sleep science. Our comprehension of such regulation in mammals has required recognizing distinct sleep stages. *Drosophila melanogaster* is an important genetic model system for studying sleep. Since the discovery of sleep-like states in the fly 25 years ago, the field has treated sleep as a unitary state consisting of any inactivity lasting 5 minutes or longer, despite convergent work suggesting the existence of multiple sleep states. Here, we establish that three distinct sleep states in flies can be classified based on simple inactivity duration criteria. We show that the daily initiation of these sleep states is temporally distinct, with long sleep occurring immediately following the largest daily period of wakefulness. We also report that rebound in response to mechanical sleep deprivation is present only in long sleep and comes at the expense of shorter sleep states. Deprivation-induced decreases in shorter sleep states obscure homeostatic sleep rebound, but only when sleep is measured using traditional methods. We observe distinctly timed ultradian oscillations of fly sleep states, reminiscent of mammalian sleep cycles. Our results indicate that the recognition of such sleep states will be necessary to fully realize the promise of the *Drosophila* model system for identifying conserved genetic mechanisms underlying such regulation.

## Introduction

With every rotation of the earth, we cycle from the “*unrecorded fantasies of solitary dreaming, to the collective fantasy of daily social and commercial life*,” as Arthur Winfree describes the mysterious and everlasting process of rhythms in sleep and wakefulness^1^. Humans spend approximately one-third of their lives asleep, about eight hours a day. Is all this sleep functionally the same? Indeed not. Early research in humans and other mammalian models recognized the existence of distinct stages of sleep during a full night’s rest^2^. Studies of mammalian sleep have made significant progress in understanding the physiological basis, functions, and consequences of sleep, progress that has depended critically on the identification and recognition of such sleep stages^3^. This is predominantly because sleep stages have distinct relationships to homeostatic and circadian regulation and correspond to different physiological states^4–12^. However, despite the identification of sleep stages in humans and other mammals, the basic biology of sleep homeostasis remains elusive^13^. In this context, the fruit fly is a promising genetic model to probe the anatomical, molecular, cellular, neural, and physiological basis of sleep homeostasis and its circadian regulation. Despite recent convergent work indicating the presence of distinct sleep states in the fly^14–24^, sleep in this species continues to be treated as a unitary state, and recent work from our lab suggests that a failure to recognize sleep states obscures reliable identification of homeostatic sleep regulation^21,22^.

*Drosophila melanogaster* sleeps for 70-80% of its life^22,25^. This estimate is based on the well-established 25-year-old behavioral correlate of sleep being any bout of inactivity that lasts longer than five consecutive minutes^25,26^. For the rest of this manuscript, we will refer to this definition of sleep as “standard” sleep. Previous work by independent groups has suggested that not all sleep is the same^14–24^. About a decade ago, physiological evidence suggested that flies may have distinct stages of sleep^17^. Specifically, the power of local field potentials recorded from the brain fell significantly after the fly had been inactive for 5 minutes, with longer periods of inactivity associated with even lower powers^17^. A subsequent study showed that metabolic rate is significantly reduced under extended periods of sleep, providing evidence for a deep sleep stage characterized by low metabolism^15^. More recently, studies have shown stereotypical proboscis extension and retraction as a marker of a discrete sleep stage^18^ and studies have demonstrated that the number of active neurons in the brain declines with the duration of sleep^19,20^.

Studies using statistical and computational approaches have also provided strong evidence in support of flies sleeping in stages^14,16^. For example, one study examined the dependence of sleep bout length on its likelihood of transitioning from sleep to wake and found that sleep may occur in at least two stages^16^. The other used Hidden Markov Models (HMMs) to characterize distinct sleep states in flies using behavioral activity data^14^. This study identified four states of fly sleep and wake. In addition to wakefulness, the authors identified “early wake”, “light sleep”, and “deep sleep” states^14^.

Based on the work of Daan and Borbély^11,12,27^, we recently modeled *Drosophila* sleep using a two-process model of sleep regulation^22^. The two-process model posits that a homeostatic process (process S; reflecting the pressure to sleep) builds while animals are awake and falls while they sleep. The switch between wake and sleep happens when process S hits an upper threshold, and the switch between sleep and wake happens when process S hits a lower threshold. The circadian clock regulates these thresholds, and the oscillating thresholds are driven by a process C^11,12,27,28^. We showed that this model captures the relationship between the circadian clock’s internal speed (free-running period) and total amount of standard sleep^22^. This relationship was also well captured by the model when we biased our analyses to include only longer episodes of sleep, which suggested that longer bouts of sleep were likely a better reflection of homeostatic sleep regulation than shorter bouts of sleep^22^.

In another recent study, we showed that only when long bouts of sleep were analyzed were we able to capture statistically significant and biologically meaningful homeostatic rebound in response to mechanical sleep deprivation in male flies^21^. When we deprived flies only of long bouts of sleep, there was a significant rebound in long sleep alone. Remarkably, when flies were deprived only of long sleep, the time course of standard sleep looked no different from that of their stimulus-matched controls. When using the standard definition of sleep, the complete loss of long-duration sleep was not detectable during the deprivation day. This result is consistent with the idea that long sleep is a more sensitive behavioral marker of homeostatic sleep regulation. It also suggests that changes in briefer sleep states can obscure homeostatic processes if not accounted for. A critical takeaway from this study was that treating sleep by the standard unitary criterion of five minutes or more of inactivity obscured homeostatic sleep regulation, and that this challenge can be easily overcome by recognizing a distinct, longer (and presumably deeper) sleep state^21^.

These convergent studies prompted us to re-examine the behavioral criterion for sleep and ask if simple behavioral criteria can identify distinct sleep states in the fly. While powerful and accurate, technical challenges are associated with the physiological and analytical observations previously used to identify deep sleep states. For instance, the physiological measurements are logistically challenging. Very few laboratories can use these systems to routinely gauge sleep stages in flies, and those that do cannot use such methods for high-throughput analyses. On the other hand, complex statistical tools such as distribution fitting and analytical tools such as Hidden Markov Models are computationally intensive, time-consuming, and require programming skills that are likely to deter widespread use. We therefore reasoned that the field would benefit if simple behavioral criteria could be used to identify fly sleep states.

In humans, the identification and characterization of sleep stages such as REM (Rapid Eye Movement) and SWS (Slow Wave Sleep), remain largely dependent on spectral analyses of Electroencephalogram (EEG) recordings^29^. While there have been significant efforts to identify such stages from peripheral physiology and movement using wearable sleep trackers, they are correlative and do not necessarily map to the same cyclic temporal architecture of polysomnography-derived sleep stages measured from the cortex. Therefore, there is a critical distinction to be made here between apparently differentiable states/types of sleep and the polysomnography-derived classification of sleep stages. Similarly, in the absence of high-throughput and an agreed-upon physiological classification of sleep stages, our definitions of sleep states in the fly must rely on the behavioral classification of apparent sleep states. However, because our states were defined in part based on previous work on physiology and metabolism, they may eventually be revealed to reflect true sleep stages.

In this study, we identified sleep states based on simple inactivity criteria and showed that they strongly resemble those identified using Hidden Markov Models. We then sought to determine if these states displayed the distinct homeostatic and circadian regulation expected of unique behavioral states. Our results established that failing to recognize sleep states in the fly obscures essential features of homeostatic and circadian sleep regulation. Thus, a full understanding of the molecular, genetic, and neuronal basis of sleep regulation will require the recognition of distinct sleep states in the fly, as it has in mammals. The recognition of sleep states will be necessary to fully realize the fly’s promise to reveal conserved mechanisms of homeostatic sleep regulation.

## Results

### Bout duration-based classification reveals distinct sleep states in Drosophila

Given previous work indicating the existence of distinct sleep states in *Drosophila*^14–20,23,24^ and the association of longer bouts of inactivity with a deeper sleep state with reduced metabolism^15^, we asked if we could define distinct sleep states based on simple inactivity duration criteria. We first considered the work of Stahl et al.^15^ (Fig. 1A), which examined the relationship between sleep bout duration and metabolic rate. The authors reported a statistically significant reduction in metabolic rate, compared to wakefulness, at ∼35 minutes into a bout of inactivity, suggesting the initiation of a deeper sleep state. We compared the time course of this metabolic change to studies of local field potential and neuronal activity in the brain in relation to the duration of inactivity^17,20^. These revealed a shift in physiological parameters ∼20 to 30 minutes after initiating a sleep bout. Taking these findings together, we reasoned that ∼30-min of inactivity might represent a cut-off between a lighter and deeper sleep state (Fig. 1A). Stahl et al.^15^ (Fig. 1A) also reported that the metabolic rate continues to decline beyond 35-min of inactivity, with significant decreases in metabolism continuing until ∼50-65 minutes after sleep initiation^15^ (Fig. 1A). Based on this, we set a second cut-off at ∼60 minutes of inactivity as a potential point of initiation of an even deeper sleep state (Fig. 1A).

**Figure 1:**
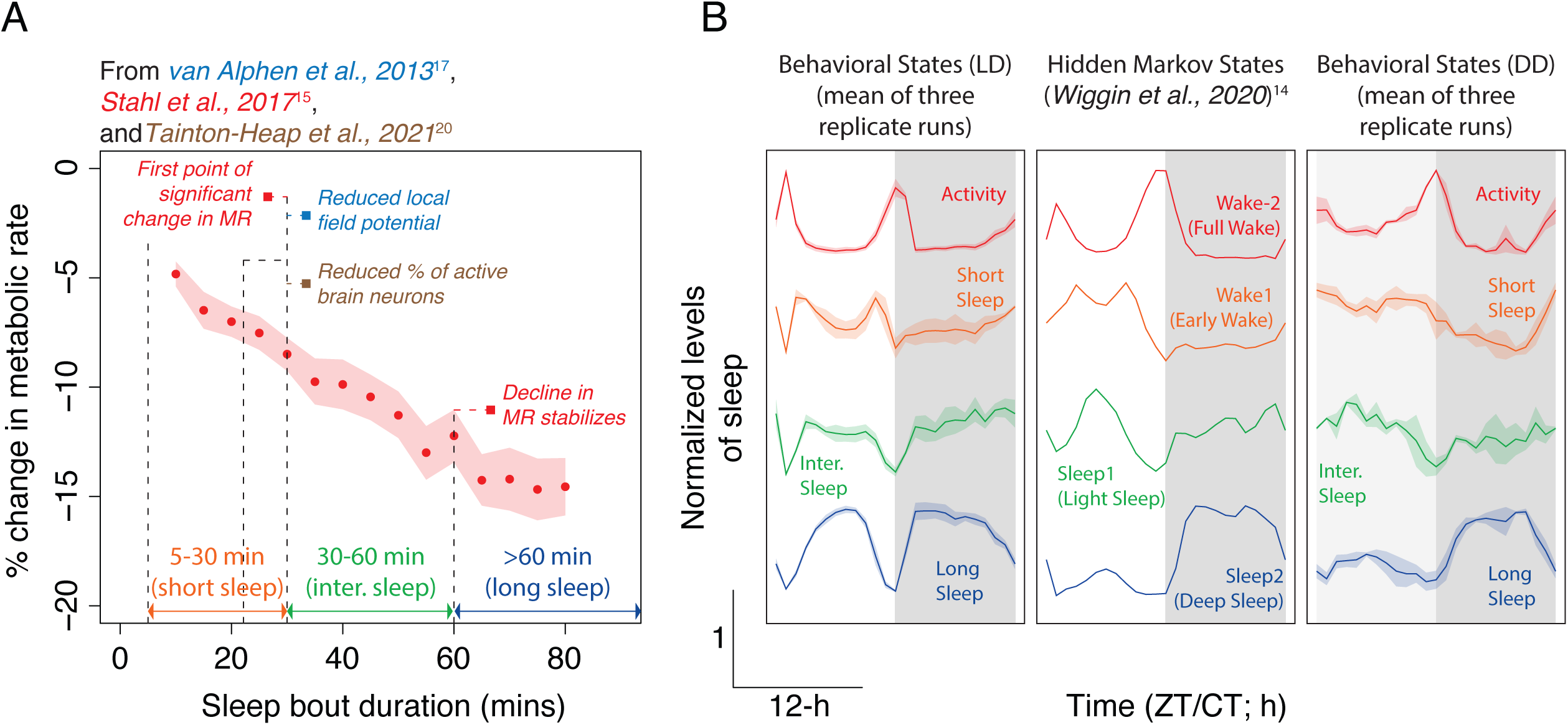
Durations of inactivity can be used to identify sleep states. (A) Plot of sleep bout durations *vs* % change in metabolic rate. Data extracted from Stahl et al., 2017^15^ and replotted here. Physiological changes previously reported by others to accompany sleep bout duration that informed bout duration cutoffs are also indicated^17,20^ (see text for details). (B) Bout length-based classification of sleep reveals waveforms of provisional sleep states under Light/Dark cycles (left) for wildtype flies. These are remarkably similar in shape to the *Drosophila* sleep states identified previously using Hidden Markov Models (HMMs; middle)^14^. Also shown are the waveforms of the three provisional sleep states, during the first day of constant darkness (right) following entrainment to LD cycles. The plots under LD (left) and DD (right) show the mean waveform from three independent replicate runs. The error regions represent the Standard Error of Mean (SEM) across the three replicate runs. Also see Suppl. Table 1.

Using these inactivity duration criteria, we provisionally defined three states of sleep, i.e., “short” sleep: 5-30 minutes inactivity, “intermediate” sleep: 30-60 minutes inactivity, and “long” sleep: > 60 minutes inactivity. But do these represent distinct states of fly sleep? To address this question, we compared the daily waveforms of these three states with a recent study by Wiggin and colleagues that used Hidden Markov Models to identify four distinct sleep/wake states, i.e., full wake, early wake, light sleep, and deep sleep^14^. First, we compared the waveforms of our provisional sleep states under Light/Dark (LD) cycles (Fig. 1B-left) with those of Wiggin et. al., whose observations were made under LD conditions (Fig. 1B-middle). Statistically significant correlations revealed that there was a remarkable similarity between the waveforms of our provisional sleep states and those inferred by the Hidden Markov Model (Fig. 1B; Suppl. Table 1). Short sleep displayed two small peaks during the daytime, much like the two peaks in early wake inferred by the Hidden Markov Model (Fig. 1B; Suppl. Table 1). Intermediate sleep was also high during the daytime, much like the light sleep state of the Hidden Markov Model (Fig. 1B; Suppl. Table 1). Both short and intermediate sleep states were low during the early part of the night and slowly increased in anticipation of dawn, as was the case in early wake and light sleep states of the Hidden Markov Model (Fig. 1B; Suppl. Table 1). Long sleep, as defined here, and deep sleep of the Hidden Markov Model also showed remarkable similarities (Fig. 1B; Suppl. Table 1). They were both high immediately after the onset of darkness and gradually ramped down in anticipation of dawn (Fig. 1B; Suppl. Table 1). However, there were small differences in the levels of our daytime long sleep and the HMM’s deep sleep (compare Fig. 1B-left and middle). While the Hidden Markov Model revealed low daytime and high nighttime levels of deep sleep, we found long sleep to be high during both day and night times. Such differences may be caused by differences in light intensities between our studies, which may cause different levels of long/deep sleep, as suggested previously^30^. Because sleep rhythms are endogenous, we asked if the provisionally defined sleep states had similar waveforms under constant darkness (DD; Fig. 1B-right). We found that the waveforms of the sleep states were largely similar to those observed under LD cycles. Interestingly, the long sleep waveform resembled the deep sleep stage inferred by the HMMs more closely under DD than under LD (Fig. 1B; Suppl. Table 1), suggesting that levels of sleep states may be differentially sensitive to the light environment.

We take these similarities as strong convergent evidence for multiple sleep states in the fly. Notably, the temporal waveforms of sleep states being largely similar under constant darkness and LD cycles indicated their endogenous nature. The distinct temporal waveforms of sleep states suggest that these states have different relationships to the major bouts of wakefulness and one another.

### The probability of initiation of long sleep is highest immediately following the largest bout of wakefulness

Our recent work indicated that long sleep is a more sensitive reflection of homeostatic sleep than briefer stages of sleep^21,22^. If true, this would predict that long sleep will be more likely than short and intermediate sleep states to occur immediately following the day’s largest bouts of wakefulness. This expectation of long sleep having a close temporal relationship with locomotor activity is also based on mammalian sleep studies establishing that different stages of sleep have distinct relationships to the sleep homeostat^11,12^.

To test this prediction, we analyzed temporal patterns of the three sleep states under various environmental conditions and compared these to the standard sleep measurements traditionally employed by the field. We examined patterns of locomotor activity and sleep and the probability of initiation of sleep bouts of each of our provisional states under (i) LD12:12, (ii) DD, and (iii) UV- and blue-blocked ramped light cycles consisting of 12-h of continuously increasing light intensity followed by 12-h of continuously decreasing light intensity. Most previous work on fly sleep has employed LD cycles. We examined sleep under DD to observe endogenous rhythms in the three sleep states, free from the acute effects of LD cycles on sleep. In addition, we examined rhythms in the sleep states under a light entrainment condition that does not produce daily startle responses, to study the entrained patterns of sleep that are not disrupted by light-induced masking effects or arousal. We chose to use UV- and Blue-blocked light cycles based on our recent work establishing that wildtype flies entrain robustly and have no startle responses of locomotor activity to any transition in light cues under these conditions^31^ and because flies actively avoid blue light during most of the day^32^.

If the homeostat regulates a particular state of sleep more closely than others, we expect that state of sleep to immediately follow the major periods of wakefulness within a diurnal or circadian cycle. We estimated the proportion of short, intermediate, and long sleep states initiated within every 1-h window across the day (see Methods). We refer to this metric as the probability of initiation of sleep bouts. In LD cycles, we found that the probability of initiating standard sleep was not significantly elevated following the most prominent daily peak of wakefulness (i.e., the evening peak of activity; Fig. 2A; Suppl. Tables 2 and 6). Thus, when measured using the standard unitary definition of sleep, it lacked a defining hallmark of homeostatically controlled rest. The probability of initiating short sleep was highest during evening anticipatory activity, which was higher than the probability of initiating short sleep following the evening activity peak (Fig. 2A; Suppl. Tables 3 and 6). Intermediate sleep initiation was most likely to occur in the hours preceding dawn and was not strongly rhythmic (Fig. 2A; Suppl. Table 4). Moreover, the probability of initiating intermediate sleep was not different before and after the evening peak of activity (Fig. 2A; Suppl. Table 6). In stark contrast, the probability of initiating long sleep was highest in the hour immediately following the evening peak of activity and was significantly higher than the probability of entering long sleep in the hours before the evening peak (Fig. 2A; Suppl. Tables 5 and 6). The probability of entering all three states of sleep displayed small increases following the relatively brief daily morning peak of activity (Fig. 2A; Suppl. Tables 2-6). Might a homeostat engage all three states of sleep during morning wakefulness, or are the small peaks in initiation probabilities following the morning peak the result of light-mediated startle effects? Is the daily peak in the probability of long sleep initiation simply a result of the rapid transition to darkness? To test this, we examined the patterns of initiation likelihood of these sleep states under DD and UV- and Blue-blocked light cycles, which do not produce the large, light-induced startle responses observed under LD.

**Figure 2:**
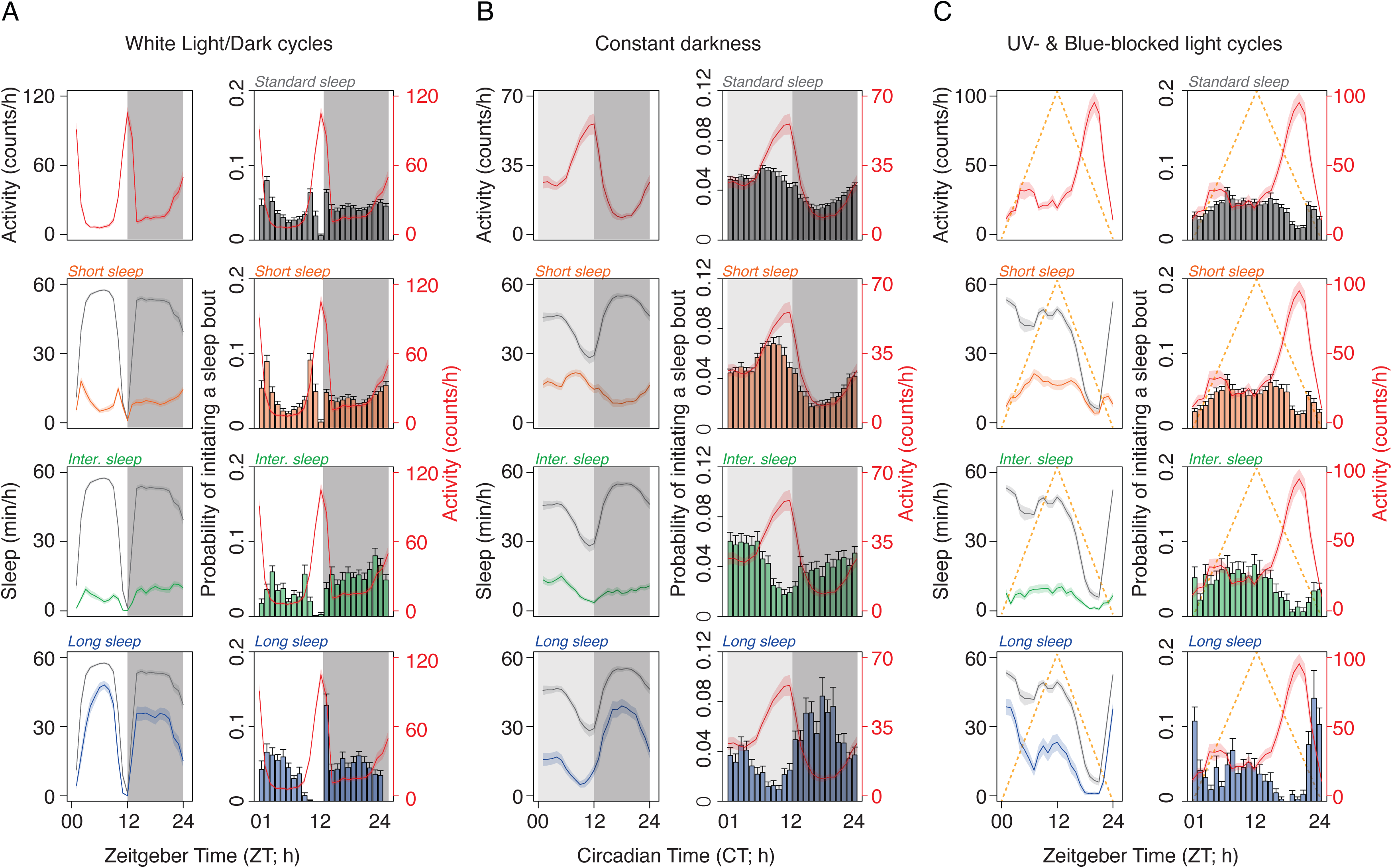
Probability of initiating long sleep is highest immediately following major peaks of wakefulness. (A) Activity and profiles of standard sleep and the three provisionally identified sleep states for wildtype flies under Light/Dark cycles (A-left), constant darkness (B-left), and UV- and Blue-blocked ramped light cycles (C-left). Also shown are plots of initiation probabilities of standard, short, intermediate, and long sleep under Light/Dark cycles (A-right), constant darkness (B-right), and UV- and Blue-blocked ramped light cycles (C-right). Gray shaded region in panel A represents the nighttime of the Light/Dark cycle. Light and dark gray shaded regions in panel B represent the subjective day and night times, respectively. The golden dotted line in panel C represents the ramping up and down of the light cycle. In the left side of all panels, the gray line represents profiles of standard sleep. These have been plotted along with profiles of the three sleep states to facilitate comparisons. Also note that the activity counts remain the same within each column, irrespective of the sleep state being represented. All the plots show mean±SEM. Also see Suppl. Tables 2-16.

Under DD, we observe a low-amplitude rhythm in the probability of initiating standard sleep (Fig. 2B; Suppl. Table 7), which peaked around mid-subjective day, when activity was increasing. The initiation of standard sleep was lowest during the early subjective night in the hours following the peak of activity (Fig. 2B; Suppl. Table 11). Short sleep was most likely to be initiated a few hours before the activity peak and was least likely to be initiated in the hours following activity (Fig. 2B; Suppl. Tables 8 and 11). The likelihood of intermediate sleep initiation was lowest during peak activity but highest near the beginning of the subjective day, in the hours preceding the largest peak of activity (Fig. 2B; Suppl. Tables 9 and 11). In contrast, the initiation of long sleep was most probable in the hours immediately following the peak of activity and was lowest during times when evening activity was ramping up (Fig. 2B; Suppl. Tables 10 and 11). These results strongly support the hypothesis that long sleep represents a sleep state potently controlled by the fly’s sleep homeostat, whereas the short and intermediate sleep states are not. Furthermore, an examination of standard sleep revealed that it was highest during times within the circadian cycle when locomotor activity was increasing (Fig. 2B). This is not how homeostatically regulated sleep is predicted to unfold. These results suggested that employing the standard, unitary definition of sleep obscured its homeostatic control during regular sleep-wake cycles.

Similar to our results in DD, the probability of initiating standard sleep had a low amplitude rhythm under UV- and Blue-blocked light cycles (Fig. 2C; Suppl. Table 12). The likelihood of initiating standard sleep was similar before and after the major locomotor activity peak (Suppl. Table 16). This was also true for short sleep (Suppl. Tables 13 and 16). Moreover, the probability of initiating short sleep was similarly high throughout the day preceding the ramping up of locomotor activity (Fig. 2C; Suppl. Tables 13 and 16). Intermediate sleep also had very weak rhythms in its likelihood of initiation (Suppl. Table 14). The highest probability of initiation of intermediate sleep was around midday, more than 12 hours after the peak of locomotor activity (Fig. 2C; Suppl. Table 16).

On the other hand, long sleep showed a much higher amplitude rhythm in the likelihood of its initiation (Suppl. Tables 15 and 16). Locomotor activity under UV- and Blue-blocked light cycles was not unimodal (Fig. 2C). There was a slight morning peak followed by a much larger evening peak. Interestingly, there was a proportionate trend in the likelihood of initiating long sleep immediately following the small morning peak (Fig. 2C; Suppl. Table 16). The probability of initiating long sleep was much higher immediately after the major peak of locomotor activity (Fig. 2C; Suppl. Table 16). These results indicate that the likelihood of initiating long sleep is directly and proportionally related to amounts and time of wakefulness and, therefore, is a closer marker of homeostatically regulated sleep. Moreover, it is imperative to note that light environments drastically affect the temporal patterns of the likelihood of initiating sleep and, therefore, must be accounted for when sleep homeostasis is being examined. Indeed, studies in flies have shown effects of light intensity and wavelength on sleep levels^30,33^. More recently, studies in rats have also shown that the wavelength of light can differentially affect various stages of sleep^34^.

### Sleep states occur in a specific temporal order relative to one another

Our results revealed that the likelihood of initiating behaviorally defined sleep states had a specific temporal order within a day. Furthermore, they indicated that long sleep displayed the relationship to wakefulness expected of a sleep state under strong homeostatic regulation. We therefore asked if these sleep states are gated to occur at characteristic times of the diurnal/circadian cycle. In mammals and other organisms in which sleep stages have been identified, the stages of sleep have a specific temporal relationship to each other, with a greater proportion of time spent in non-rapid eye movement (NREM) and REM sleep during the earlier and later parts of the night, respectively^35^. A critical goal of sleep science is to understand the temporal architecture of these sleep stages and how they are coordinated by the brain^3^. Because of the large startle/arousal-mediated effects on sleep states under LD cycles (Fig. 2A), we examined the temporal organization of sleep states only under DD and UV-and Blue-blocked ramping light cycles, which do not produce such acute effects on activity.

As expected, most of the locomotor activity is concentrated around subjective dusk, under DD conditions (Fig. 3A; subjective dusk corresponds to the time of lights-off under the LD cycle preceding transition to DD, and subjective dawn corresponds to lights-on). Furthermore, standard sleep was spread across nearly 20 hours of the circadian cycle, corresponding to all times outside of the subjective evening peak of activity (Fig. 3A). Short sleep occurred primarily within the subjective day and peaked around three hours before subjective dusk, just before locomotor activity started ramping up (Fig. 3A). Intermediate sleep also occurred primarily within the subjective day but occurred mainly in the first half of the subjective day (Fig. 3A). Long sleep was predominantly timed to the subjective night and started building up soon after the peak of locomotor activity (Fig. 3A). Thus, the timing and amount of the three sleep states matched the probability of initiation described above (Fig. 2).

**Figure 3:**
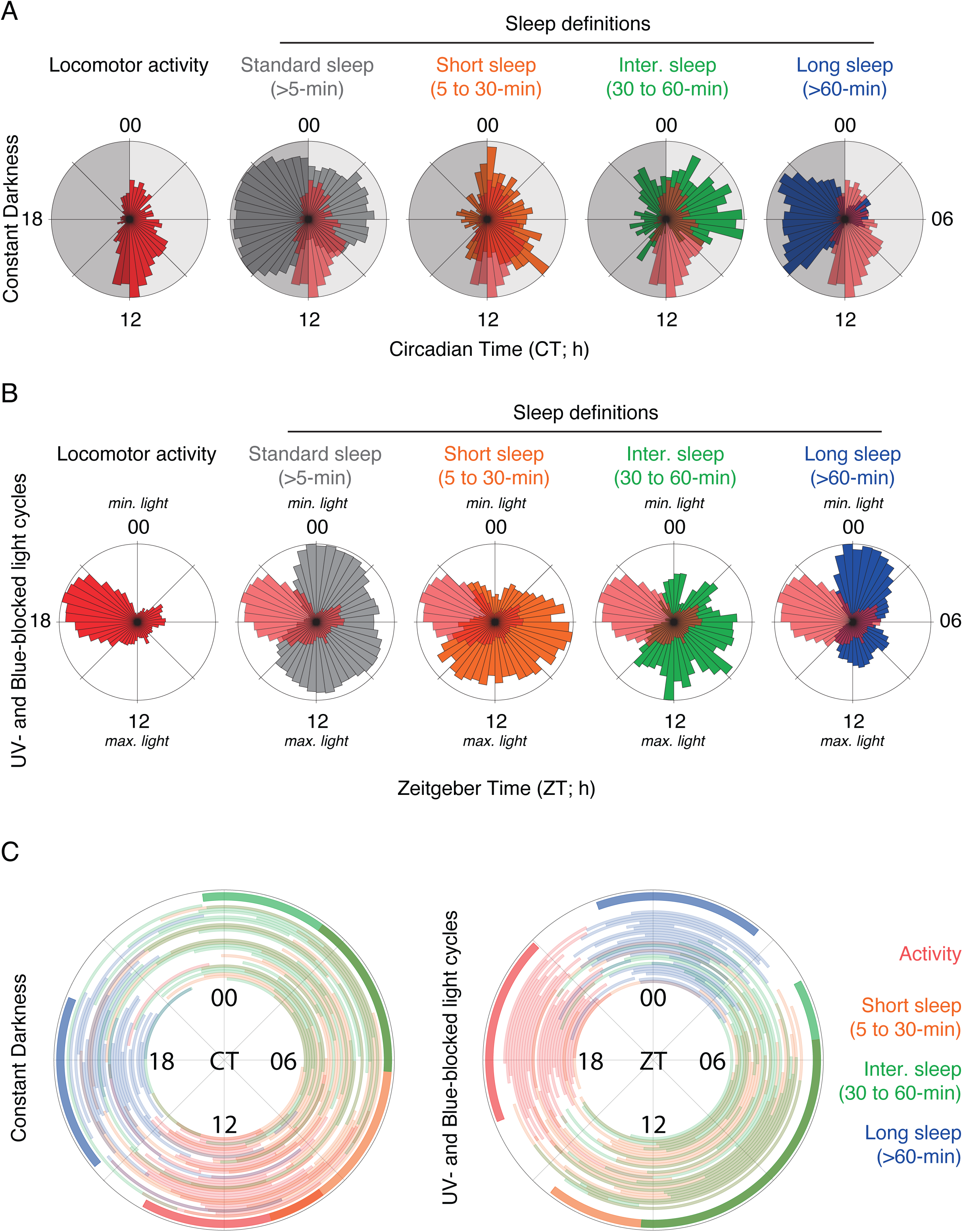
Rose plots clearly demonstrate the temporal sequencing of sleep states in wildtype flies. Rose plots of locomotor activity and sleep states under constant darkness (A) and UV- and Blue-blocked light cycles (B). Rose plots are averaged over flies. In each sleep state, the locomotor activity rose plots are overlaid to facilitate visualization of their temporal inter-relationships. (C) Polar plots representing the gating of the four sleep/wake states under constant darkness (left) and UV- and Blue-blocked light cycles (right). Each concentric ring shows the gate of each sleep/wake state for each fly. The outermost ring with thicker arcs represents the mean circadian gating of each sleep state. These analyses allowed us to extract onset and offset phases of each sleep state, thereby providing a quantitative description of the temporal organization of these sleep states. Note that although sleep states cannot co-occur in time within a fly, the circadian gates may overlap. See Suppl. Fig. 4 for an example. Also see Suppl. Tables 17-18.

When entrained, under UV- and Blue-blocked light cycles, fly sleep states displayed a temporal sequencing similar to that observed under constant darkness. Rose plots revealed that long sleep predominantly occurred in the hours closely following locomotor activity, which was centered on the falling phase of the ramping light cycle (Fig. 3B). There was a slight, proportionate increase in long sleep closely following the minor peak of locomotor activity observed during the falling phase of the light ramp (Fig. 3B). Most of the short and intermediate sleep states occurred within broad windows preceding the major bout of locomotor activity (Fig. 3B), similar to their timing under constant darkness.

Statistical comparisons of such temporal sequencing of sleep states require the ability to estimate the phases of onset (*ψ_onset_*) and offset (*ψ_offset_*) of each sleep state. However, due to the lack of objective assessments of phase markers to identify the onset and offset of sleep states, we first derived a proxy to estimate their timing (see Methods). It was clear that long sleep was gated immediately following locomotor activity and the short and intermediate sleep states were gated to a window following long sleep, just prior to the onset of locomotor activity, under both environmental conditions (Fig. 3C). To test this statistically, we carried out the following planned comparisons: (i) activity *ψ_offset_ vs* long sleep *ψ_onset_*, (ii) long sleep *ψ_offset_ vs* short sleep *ψ_onset_*, (iii) long sleep *ψ_offset_ vs* intermediate sleep *ψ_onset_*, (iv) short sleep *ψ_offset_ vs* activity *ψ_onset_*, and (v) intermediate sleep *ψ_offset_ vs* activity *ψ_onset_*. Under both constant darkness and UV- and Blue-blocked light cycles, we found that the *ψ_onset_* of long sleep occurred significantly later compared to the *ψ_onset_* of activity (Fig. 3C; Suppl. Tables 17-18). Furthermore, the *ψ_onset_* of short sleep occurred significantly after the *ψ_offset_* of long sleep (Fig. 3C; Suppl. Tables 17-18). Under constant darkness, the *ψ_onset_* of short intermediate sleep also occurred significantly after the *ψ_offset_* of long sleep, whereas such a statistically significant difference was not detectable under UV- and Blue-blocked light cycles (Fig. 3C; Suppl. Tables 17-18). When we compared the *ψ_offset_* of short and intermediate sleep against the *ψ_onset_* of activity, we found that, under UV- and Blue-blocked light cycles, the gate of activity opened after the gate of short sleep closed (Fig. 3C; Suppl. Table 18). Although a similar trend was seen under constant darkness, statistical significance was not attained (Fig. 3C; Suppl. Table 17). Under both environmental conditions, the gate of activity opened after the gate of intermediate sleep closed (Fig. 3C; Suppl. Tables 17-18).

The temporal segregation between short and intermediate sleep states was less drastic under ramped light cycles than under constant darkness. This may have been due to the small secondary peak of locomotor activity during the rising phase of the light cycle (Figs. 2C and 3B), which may have resulted in shorter sleep states being distributed more widely around the clock. Nonetheless, there was a clear temporal sequencing of sleep states under both constant darkness and entrainment to ramped light cycles: long sleep followed locomotor activity, which was followed by intermediate sleep, and subsequently by short sleep. The consistent daily timing of the long sleep state to the hours following peak wakefulness further supports the hypothesis that the sleep homeostat more potently controls this sleep state than the short and intermediate states of sleep.

### The homeostatic rebound of long sleep occurs at the expense of shorter sleep states, obscuring rebound measures when the standard, unitary definition of sleep is used

Long sleep was unique among the sleep states in its tendency to follow the day’s major bout of wakefulness, an expected hallmark of a sleep state regulated by homeostatic mechanisms. One of the most direct ways of examining homeostasis is to examine sleep after sleep deprivation^25,26^. We further explored the relationship between our provisional sleep states and homeostatic regulation by asking if rebound following sleep deprivation produces differential effects on short, intermediate, and long sleep states. Existing approaches to sleep deprivation in the fly produce relatively modest rebounds^21,36,37^. Using the Ethoscope system^38^, we recently showed that homeostatic sleep rebound was obscured when sleep was defined using the standard approach as any bout of inactivity lasting five minutes or longer^21^. However, when more prolonged sleep bouts were examined, in this case, defined as periods of inactivity lasting 25 minutes (owing to video-based measurements; see Discussion) or longer, a significant proportion of lost sleep was recovered following deprivation. Interestingly, when flies were completely deprived of only long sleep, the time courses of standard sleep (five minutes of inactivity or longer) looked no different from those of their stimulus-matched controls^21^. Nevertheless, analysis of long sleep revealed a significant cumulative rebound over multiple cycles following sleep deprivation^21^.

Though Ethoscopes are a powerful and flexible method of depriving flies of sleep, they have not yet been adopted widely by the field. A more commonly used method of sleep deprivation is the vortexer-induced mechanical disturbance combined with Trikinetics *Drosophila* Activity Monitors to measure sleep through beam crossing measurements^39^. This method has consistently produced, in many different labs, a measurable, albeit modest, sleep rebound following deprivation when flies are stimulated three times per minute to prevent sleep. We recently showed that such mechanical disturbances delivered less frequently prevent sleep completely yet produce no statistically significant rebound in standard sleep^21^. These results prompted us to ask if standard sleep measurements obscured homeostatic sleep rebound when using the vortexer method of sleep deprivation. In mammalian systems, sleep stages are characterized by differential responses to deprivation^2,40^. We therefore asked if the recognition of sleep states in the fly might reveal hidden homeostatic responses. Distinct responses to sleep deprivation would be strong evidence that our provisional sleep states represent distinct behavioral states.

Despite being an important and frequently used experimental approach, methods for depriving flies of sleep and measuring sleep rebound vary significantly between studies and have not been standardized^13,39^. This has made it difficult to compare reported effects on homeostatic sleep regulation between studies^13^. Owing to the strong circadian regulation of sleep, we recently suggested the use of 24-h sleep deprivation and a 24-h rebound analysis window^21^, as opposed to a large number of previously published studies depriving for shorter periods and examining sleep rebound within a smaller window of time following deprivation^39^. Here we employ a simple method for analyzing homeostatic sleep rebound after sleep deprivation (Fig. 4A). We define sleep rebound as the difference in sleep amount between the post deprivation period and baseline day, adjusted for the changes in sleep displayed by undisturbed controls across the three days during which baseline, deprivation, and recovery sleep are measured (Fig. 4A; see Methods).

**Figure 4:**
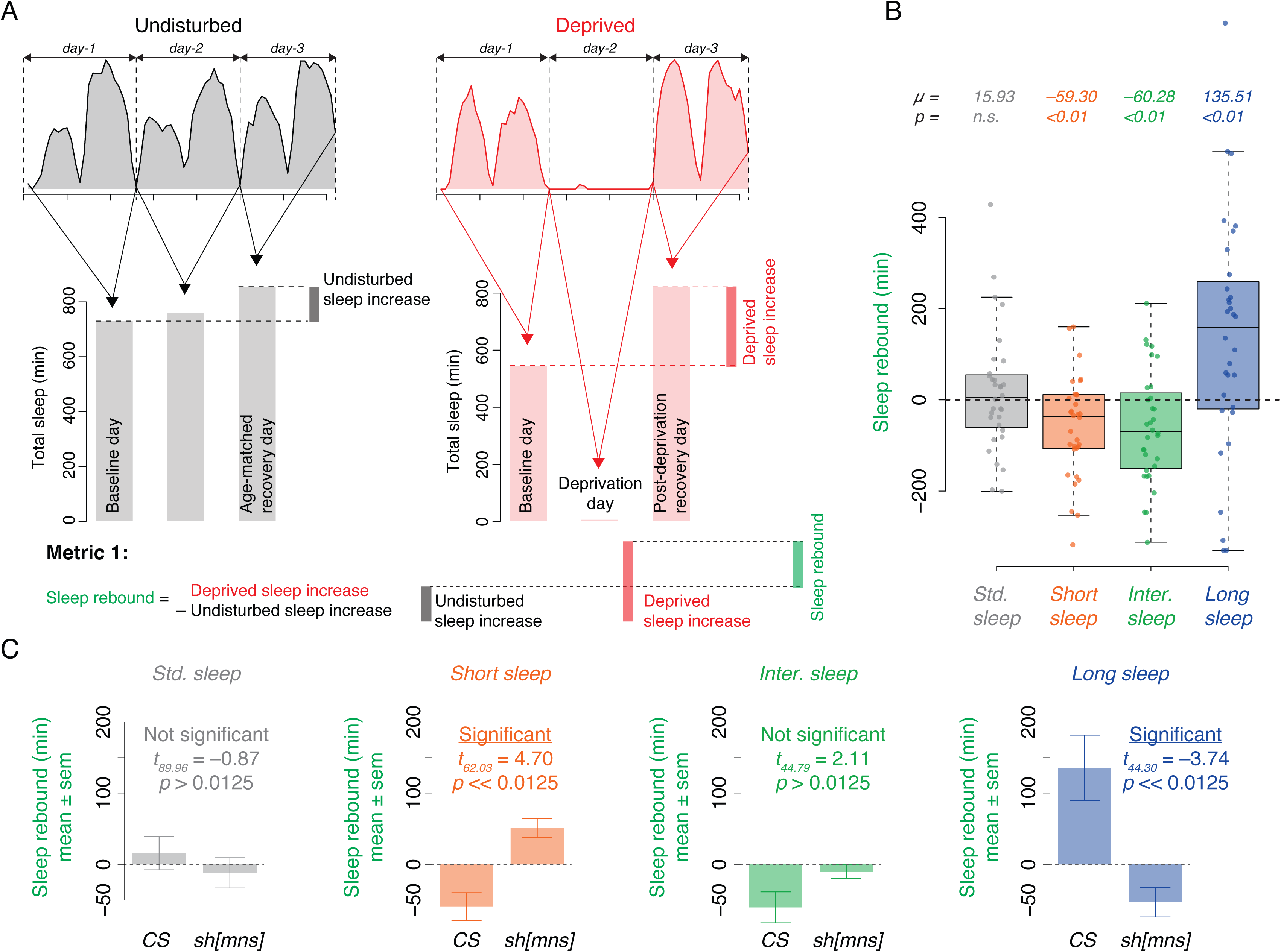
Long sleep displays substantial homeostatic sleep rebound following deprivation, rebound that is obscured by standard sleep, in wildtype flies. (A) Schematic of a proposed method of measuring of sleep rebound following deprivation. (B) Estimated sleep rebound for standard sleep and each of the three provisional states of sleep. Statistical comparisons were made against 0 for each of the metrices and sleep stages separately using single sample *t*-tests. (C) Estimated sleep rebound for standard sleep and each of the three provisional sleep states for a “homeostatic” mutant and its background control, *CS* flies. Note that the data for CS flies is replotted from panel B. Data for the *sh[mns]* are pooled from five independent runs (n = 156). Also see Suppl. Figs. 1-2.

We calculated sleep rebound for each of our provisional sleep states to determine if the sleep states proposed here respond differentially to sleep deprivation. As we previously reported^21^, we found no statistically significant rebound in standard sleep over 24-h post deprivation when male flies were kept awake for 24 hours by mechanically vortexing them once every 220-sec (Fig. 4B; Suppl. Fig. 1; note that the field has typically stimulated flies every 20-sec to prevent sleep). In contrast, there was a significant rebound in long sleep (Fig. 4B; Suppl. Fig. 1). Flies showed more than two hours of rebound in long sleep (Fig. 4B; Suppl. Fig. 2). Remarkably, this increased rebound in long sleep occurred at the expense of short and intermediate sleep states, both of which showed statistically significant reductions following deprivation (Fig. 4B; Suppl. Fig. 1). These results establish that our provisional sleep states display distinct responses to sleep deprivation, suggesting that they do represent different behavioral states. Though sleep states are, by definition, mutually exclusive, this does not mean that changes in one state must occur at the expense of others if the total amount of sleep changes. Our results reveal a distinct homeostatic regulation of the sleep states defined here and demonstrate the necessity of distinguishing sleep states when analyzing the homeostatic regulation of fly sleep.

To further illustrate the utility of our method of examining sleep homeostasis, we measured levels of sleep in response to deprivation in a previously characterized “homeostatic” mutant, i.e., *shaker[minisleep]* (*sh[mns]*)^41^. We used *Canton-S (CS)* as a background control for *sh[mns]* in our comparisons owing to the fact that *minisleep* flies were generated in a *CS* background^41^. The *minisleep* mutation in *shaker* was previously reported to cause a decrease in baseline sleep on multiple genetic backgrounds but did not appear to prevent homeostatic rebound following deprivation^41^. For standard sleep measurements, we found that there was no significant difference in rebound between *CS* and *sh[mns]* flies, as was reported previously^41^ (Fig. 4C; Suppl. Fig. 2). However, there was a significant increase in short sleep rebound and a significant decrease in long sleep rebound in *sh[mns]* flies compared to *CS* flies (Fig. 4C; Suppl. Fig. 2). We did not detect any significant difference between the genotypes in rebound for intermediate sleep (Fig. 4C; Suppl. Fig. 2). Our analyses strongly suggest that *shaker* flies indeed have a considerable rebound phenotype, but only when sleep states are recognized – further highlighting the fact that standard definitions of sleep may obscure homeostatic effects. We confirmed the mutant line by comparing their baseline sleep under LD, which showed approximately similar reductions in sleep compared to the background controls as previously reported (>48% reduction)^41^. All our rebound results emphasize a critical point: treating sleep as a unitary state obscures readouts of homeostatic regulation.

### Long sleep displays robust circadian rhythms, whereas short and intermediate sleep states display stronger ultradian rhythms

Distinct sleep states likely have different functions and may, for that reason, be differentially controlled by the circadian system. Previous work on mammalian sleep has shown that REM (rapid eye movement) and NREM (non-REM) sleep have distinct relationships to the circadian clock. While the central circadian system sets the timing of both these sleep stages, and NREM and REM are both homeostatically regulated, the circadian clock appears to have a stronger influence on REM sleep^4–12^. Though the circadian clock gates mammalian sleep, once sleep commences, alternations between sleep stages result in ultradian rhythms of sleep as it cycles through multiple stages. Thus, circadian rhythms in sleep are accompanied by ultradian rhythms in specific sleep stages during particular phases of the circadian cycle. Therefore, we examined both circadian and ultradian periodicities to determine whether the three provisional sleep states we identified exhibit distinct circadian regulation. We employed wavelet analyses to resolve periodic components of sleep time series across days under constant darkness, as we have recently described^22^. Wavelet-based methods of timeseries analyses allow for the identification of periodic components in the timeseries and how they unfold over time^42–44^.

Scalograms visualize the power of the rhythm for each periodic component of a timeseries across time (see Methods). The power of a particular periodicity reflects the amplitude of the oscillation. The terms rhythmic power, strength, and amplitude are used interchangeably here. Scalograms of standard sleep revealed strong 12- and 24-h periodicities, as expected from the fly’s biphasic sleep pattern (Fig. 5A). In addition, there were strong ultradian periodicities present under DD, whose predominant periods were in the one to four-hour range (Fig. 5A). These ultradian periodicities largely displayed relatively low amplitudes compared to the 12-and 24-h rhythms in standard sleep (Fig. 5B). We repeated this analysis for the three states of sleep that we provisionally identified above to determine if they display distinct circadian control and temporal organization under free running conditions, as would be expected of distinct sleep states.

**Figure 5:**
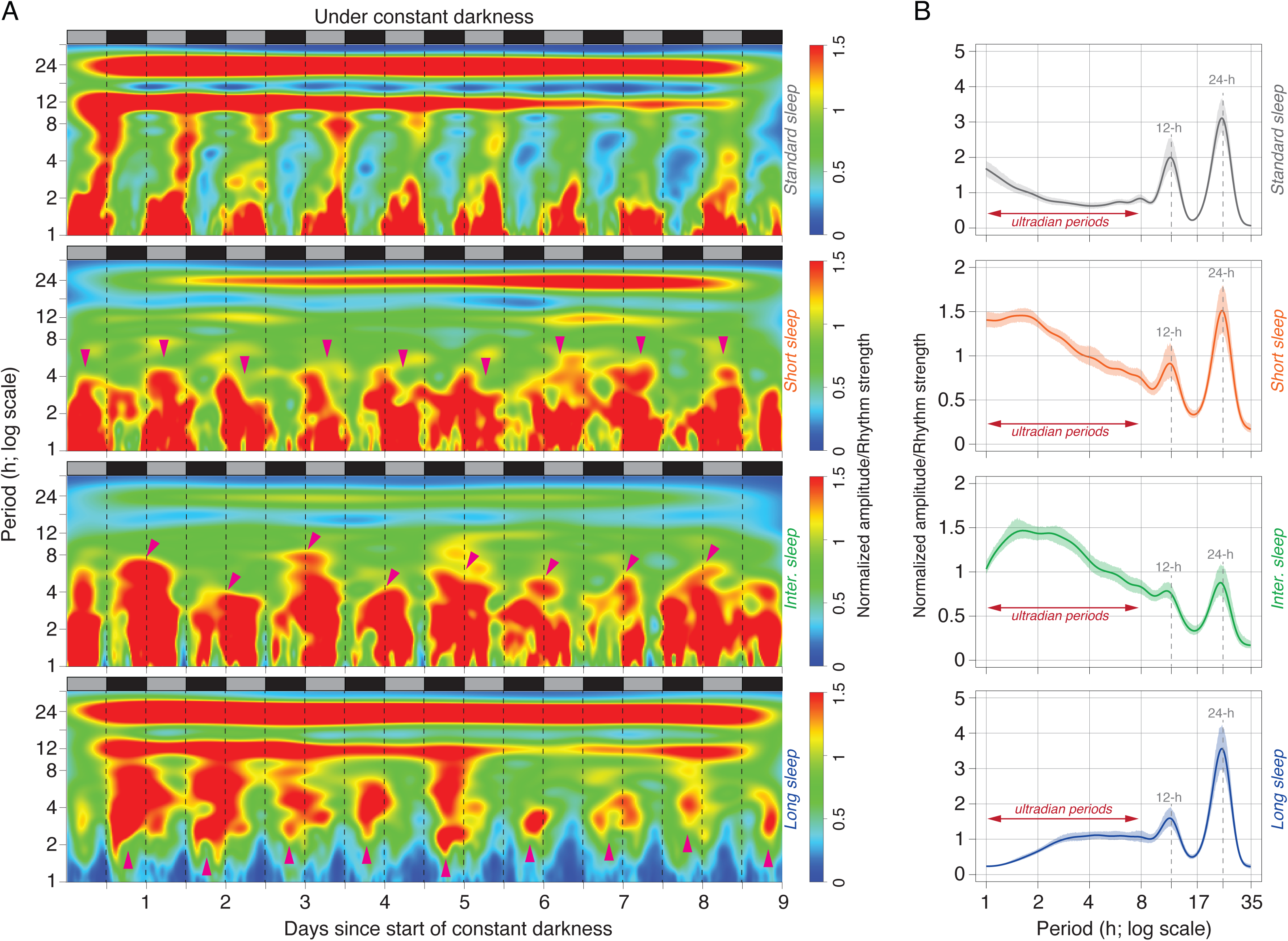
Sleep states display distinct circadian and ultradian regulation. (A) Normalized, averaged scalograms of standard sleep, and the three states of sleep under constant darkness. Note that the *y-*axis is plotted on a logarithmic scale. Magenta arrowheads are meant to indicate potential gating of ultradian rhythms as reflected by relatively high amplitudes. Light and dark gray shaded regions represent the subjective day and night times, respectively. (B) Plots of period *vs* amplitude for all the definitions of sleep under constant darkness, averaged over the time-axis. Note that *x*-axis is plotted on a logarithmic scale. The error regions for each trace represent 95% confidence intervals estimated through bootstrapping. Therefore, any amplitude values that have non-overlapping error regions may be considered statistically significantly different from each other.

An examination of short sleep revealed high amplitude cycling in both the 24-h range and ultradian periodicities with relatively low amplitude cycling at the 12-h periodicity (Fig. 5A). When averaged across time, we found that the ultradian rhythm component was comparable in amplitude to the circadian peak (Fig. 5B). In contrast, intermediate sleep displayed relatively low amplitude periodicities in the circadian range and strong periodicities in the ultradian range (Figs. 5A and B). Finally, and in contrast to short and intermediate sleep states, long sleep displayed strong periodicities in circadian range, moderately strong periodicity in the 12-h range, and relatively low amplitude ultradian periodicities (Fig. 5A and B). These results revealed a distinct regulation of sleep states by the circadian clock and further support the need to recognize sleep states in the fly.

### The circadian clock sets the phase of ultradian sleep rhythms

In mammals, sleep cycles through various stages during the night with an ultradian periodicity. For instance, such ultradian rhythms have a periodicity of ∼90 minutes in humans^3^. An analysis of a long timeseries of a single stage of human sleep would produce a scalogram containing circadian and ultradian components, as we have observed for the fly (Fig. 5). The circadian component would be a consequence of the sleep stage being restricted to the night. The ultradian component would reflect the cycling in and out of that sleep stage, which would likewise be limited to the night. Therefore, mammalian sleep stages display circadian and ultradian periodic components, with the ultradian components restricted to specific times of the day^3^. An examination of the scalograms presented in Fig. 5A suggests that ultradian oscillations in short, intermediate, and long sleep states in the fly occur within distinct windows, with each state displaying a characteristic timing (magenta arrowheads in Fig. 5A). High amplitude ultradian rhythms of short and intermediate sleep states appeared to be timed to the subjective day, whereas the relatively higher amplitude of ultradian rhythms in long sleep were restricted to the subjective night (Fig. 5A). This prompted us to ask whether the circadian clock gated these ultradian rhythms to specific times of the day by comparing wildtype flies to clock loss-of-function *period^01^* (*per^01^*) mutants under constant darkness. If the circadian clock was responsible for the proper daily timing of ultradian cycling of sleep states in the fly, we would expect *per^01^*mutants to lack the consolidation of ultradian rhythms we observed in wildtype flies (Fig. 5A).

We found that the strength (i.e., amplitudes) of ultradian periodicities in short, intermediate, and long sleep were clearly rhythmic for wildtype flies but not for *per^01^*mutants (Figs. 6A, C, E). We tested the statistical significance of rhythmicity by subjecting these timeseries to chi-squared periodogram analyses^45^. We found significant rhythmicity in wildtype flies and no such rhythmicity in *per^01^* mutants for each of the three sleep states (Figs. 6B, D, F; Suppl. Table 19). Even the very few *per^01^* flies that had significant rhythmicity, displayed power values vastly lower than those of wildtype flies (Suppl. Table 19). Thus, these three sleep states display distinct modes of circadian regulation, further supporting the conclusion that they represent distinct behavioral states. Our results further highlight the fact that standard measures of sleep obscure important aspects of its regulation.

**Figure 6:**
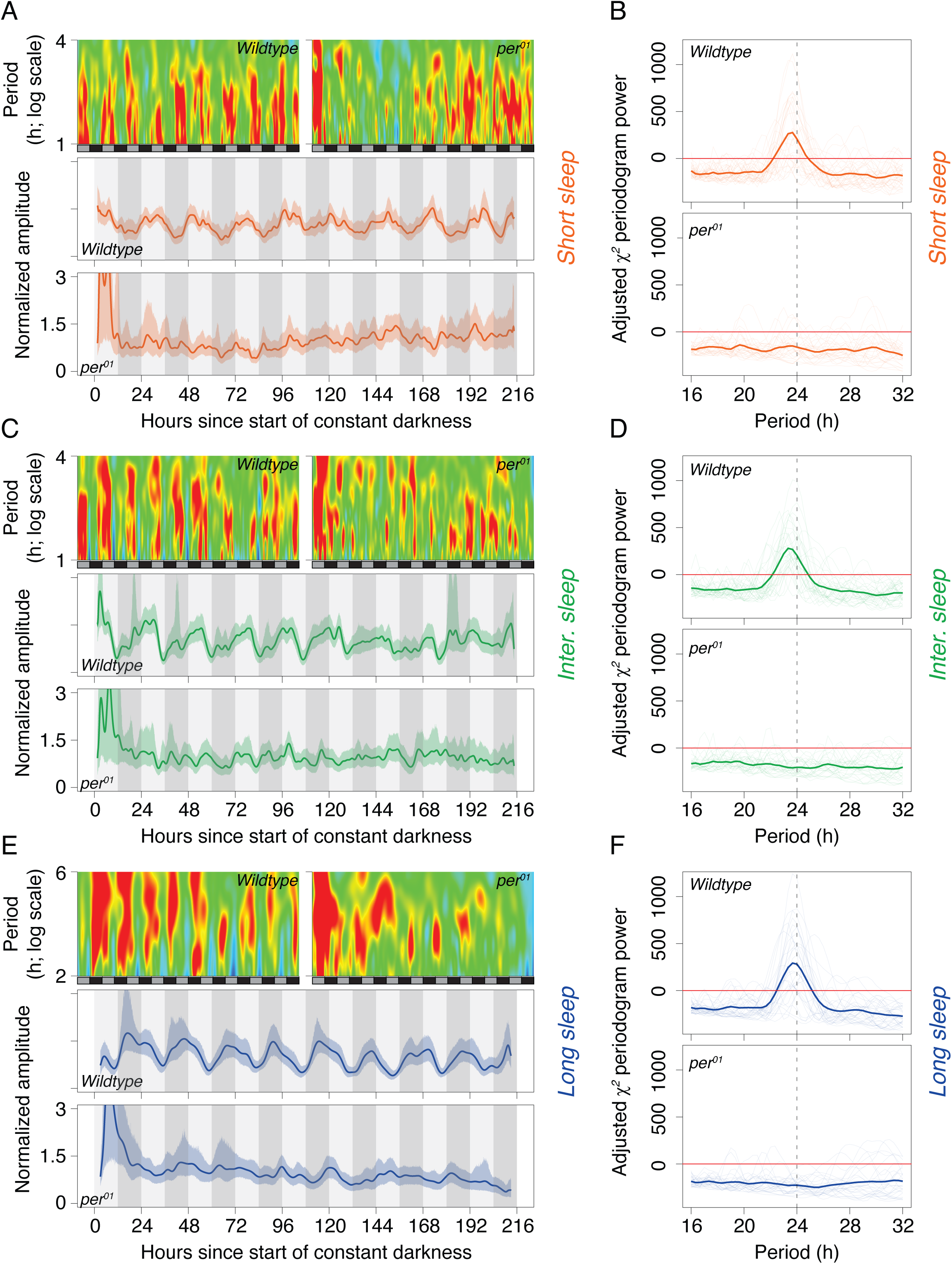
Ultradian rhythm amplitudes for each sleep state are distinctly gated by the circadian clock. Normalized scalograms for short (A), intermediate (C), and long (E) sleep in wildtype and loss-of-function *per^01^* mutant flies depicting amplitude values in the ultradian period range used for downstream analyses (see text for details). Also, shown are amplitude values over time for each sleep state (A, C, E). All scalograms have a *z-*axis scale that ranges from 0 to 1.5. The error regions for each time *vs* amplitude trace represent 95% confidence intervals estimated through bootstrapping. Results of chi-squared periodogram analyses carried out on wildtype and *per^01^* flies are plotted for short (B), intermediate (D), and long (F) sleep. Thin lines represent the periodogram results for each fly, and the thick line displays the power for each period value, averaged over all the flies. The red horizontal line at zero marks the critical value for statistical significance. Any period value with a peak above the red line is reflective of a statistically significant rhythmic component. Also see Suppl. Table 19.

## Discussion

### Do flies have distinct sleep states?

The recognition of human sleep stages was a critical discovery for sleep science, and our current understanding of sleep and its impact on physical and mental health would be significantly less advanced if human sleep had been treated as a unitary state^2,3^. Based on recent convergent work suggesting the presence of distinct sleep states *in Drosophila melanogaster*, we asked if distinct states can be defined behaviorally in this species. Indeed, we have identified four states of fly sleep/wake: wakefulness (i.e., activity), short sleep, intermediate sleep, and long sleep, which are based on previous work examining the relationships between inactivity and physiology^14–24^ and correspond remarkably well to the sleep/wake states previously identified using Hidden Markov Modeling (Fig. 1B). The states we have identified here have distinct relationships to both circadian and homeostatic regulators, as would be expected of discrete sleep states. We have also established simple inactivity-based criteria that can be used to identify these sleep states (Fig. 1).

Results described here build upon our recent work^21,22^, and show that long sleep, among the three sleep states, is uniquely subject to homeostatic control (Figs. 2-4). Importantly, long sleep shows a clear relationship to homeostatic processes during both daily sleep rhythms and rebound after sleep deprivation. Previous work using the standard unitary definition of sleep has observed an apparent disconnect between the regulation of daily sleep and homeostatic rebound based on manipulations that affect one but not the other^13^. Based on these results, investigators have suggested that baseline sleep and homeostatic rebound are governed by different molecular/cellular mechanisms. However, the two-process model of sleep regulation, which we recently showed captures fly sleep quite well^22^, posits that the process that produces sleepiness during normal wakefulness is the same process engaged during sleep deprivation. Here we show that the long sleep state, uniquely among the states identified in the study, maintains both the predicted temporal relationship to wakefulness and responds to sleep deprivation with substantial rebound (Figs. 2-4). Remarkably, the first sleep mutant identified using forward genetics revealed decreased baseline sleep, but no clear effect on homeostatic rebound^41^. Here, we have shown that this mutant displays a robust phenotype when sleep stages are accounted for. Our identification of a sleep state potently regulated by the sleep homeostat that has been obscured by the field’s unitary definition of sleep calls for the reassessment of previous work identifying genetic and cellular components of the fly’s sleep homeostat.

Our work also establishes that fly sleep comprises identifiable states that exhibit unique temporal sequencing across the circadian cycle (Figs. 2-3), as would be expected of distinct sleep stages. Moreover, we found that all three sleep states display both circadian and ultradian rhythms (Fig. 5), and that ultradian rhythms of each state are gated by the clock to specific windows within the circadian cycle (Fig. 6). Future work will be required to determine if these gated ultradian rhythms represent something akin to programmed sleep cycles in the fly.

One potential objection to our definition of sleep states in the fly is that it is simply a variation on the field’s longstanding practice of examining sleep bout duration as a measure of sleep consolidation. Sleep consisting of fewer, but longer bouts of sleep, is said to be more consolidated than the same amount of sleep divided into shorter and more frequent bouts. One might argue that the use of inactivity criteria to define distinct sleep states is simply another way of measuring consolidation. But simple arithmetic can show that the amount of long sleep can be identical in two flies with significantly different average sleep bout durations (Suppl. Fig. 3). For example, changes in the relative amounts of short and intermediate sleep can produce changes in average sleep bout duration when no changes have occurred in long sleep (e.g., Suppl. Fig. 3).

One important caveat of our provisional definition of sleep states is the data acquisition method used to examine sleep. Our definition of short (5 to 30 minutes), intermediate (30 to 60 minutes), and long (> 60 minutes) sleep is applicable only to data collected using the IR beam-based acquisition method through the Trikinetics *Drosophila* Activity Monitoring systems (Waltham, MA, USA). Compared to video-based measurements, the DAM system overestimates sleep^21,38,46,47^, and therefore, the corresponding inactivity criteria will be lower for video-based data acquisition methods. Careful calibration will be required to standardize criteria across different systems. However, based on our previous work^21^ we suggest that ∼30 minutes of inactivity in a video-based data acquisition method likely corresponds to the long sleep we define here. Remarkably, a founding study in the field of *Drosophila* sleep science, which used a video-based behavioral assay, suggested that a large percentage of all sleep was > 30-min and the longest bouts of sleep occurred soon after subjective dusk^25^. In hindsight, a 30-min inactivity criterion for sleep, based on one of the earliest reports of fly sleep^25^, our recent work^21,22^, and the results presented here, would likely have been a more sensitive metric for examining sleep homeostasis over the past 25 years.

### It is difficult and unwise to compare sleep stages across phyla directly

The circadian gating of ultradian rhythms in sleep states that we reported is reminiscent of the temporal patterning of sleep cycles in mammalian sleep^48^. It is important to highlight that we are not suggesting that the sleep states identified herein are akin to REM (Rapid Eye Movement)/NREM (non-REM) sleep in mammals, though there are clear similarities in two regards. First, like mammalian sleep stages, fly sleep states are distinctly sensitive to homeostatic regulation^21,22^. Second, different sleep states in the fly are distinctly timed to occur at different phases of the circadian cycle, much like sleep stages in mammals. Although REM/NREM sleep stages have been identified in rodent systems, they have been shown to differ somewhat from human REM/NREM stages, in that human NREM has substages^49^. There are also physiological differences between rodent sleep stages and human sleep stages^49^. Interestingly, while it is true that birds show REM and NREM sleep stages, they appear to differ from mammalian REM/NREM in important ways^50,51^. For instance, bird NREM sleep lacks substates and specific brain oscillations^50,51^. Reptilian sleep also has electrophysiologically distinct sleep stages, but REM and NREM-like sleep stages in the bearded dragon do not completely match REM and NREM sleep stages seen in mammals^52,53^. Therefore, while there may be similarities in sleep states between species, there are also likely to be significant differences across phyla, and directly comparing them, without understanding their function and regulation, is unwise^54^. In comparing sleep stages across phyla, we must be mindful of the “*misuse of nomenclature, and lack of openness towards potentially intriguing differences*” as warned by Rayan and colleagues^49^. As discussed above, it is not our intention to conflate the behavioral identification of sleep states and EEG-based classification of sleep stages in humans; instead, our recognition and characterization of sleep states in the fly may serve as markers to enable future investigations into the possibility of these states reflecting similar neurophysiological mechanisms as those revealed by polysomnography-derived sleep staging in mammals.

### Our results call for an end to the treatment of fly sleep as a unitary state

Inactivity-based measures of sleep correlate strongly with electroencephalograms (EEGs) and electromyograms (EMGs) and have been suggested as a first-pass behavioral criterion to examine sleep in the pursuit of high-throughput screening programs in mice^55^. The use of similar strategies in the fly, especially in the absence of easily accessible physiological correlates, can be readily used by the field in its effort to identify the neural and molecular genetic correlates of sleep regulation. For over 25 years, fly sleep research has used a unitary definition of sleep, i.e., any bout of inactivity equal to or longer than five continuous minutes. Despite half a century of sleep scoring using standards set by Rechtschaffen and Kales^29^, the classification of sleep remains an active area of research in mammalian sleep^49^. Why should the same not be true of sleep research in the fly?

Despite a quarter century of work and the identification of hundreds of “sleep genes,” the field has failed to produce a widely accepted model for sleep homeostasis^13^. Our results here, along with other recent work from our lab, converge to provide a possible explanation for this shortcoming: (i) the employment of a unitary definition of fly sleep, and (ii) the use of bright light/dark cycles with very little information on intensity or spectral composition of the light have obscured potent homeostatic sleep processes. Specifically, we have shown that using the standard definition of sleep obscures the detection of sleep rebound because manipulations used to deprive flies of sleep have opposing effects on different sleep stages (Fig. 4). Although every bout of inactivity longer than 5-min meets the behavioral criteria for sleep, they are not all homeostatically regulated, and most certainly not in the same manner. The recognition, refinement, and widespread acceptance of sleep states in the fly will be required if the *Drosophila* model’s promise for identifying conserved mechanisms of sleep homeostasis is to be realized. Realizing this promise will require the field to closely examine the sleep states most strongly subjected to homeostatic regulation. Treating all epochs of inactivity lasting five minutes or more as a unitary sleep state has likely outlived its utility for discovering the mechanisms underlying sleep regulation. This is particularly true for understanding the homeostatic regulation of sleep, which remains a primary unmet goal of sleep science.

## Conclusion

Sleep stages in humans are clearly identifiable and have predictable temporal sequencing^3,48^. They correlate with different physiological and behavioral changes^3^. Their identification has been critical to understanding the complex phenomenon of sleep. However, much remains to be discovered regarding the molecular and neural regulation of the timing and amount of sleep^13^. *Drosophila melanogaster* remains a highly promising model system for discovering conserved mechanisms of sleep regulation. Our identification of sleep states in the fly will allow for a more sensitive and accurate accounting of sleep regulation. The states we have defined here remain provisional and, in contrast to the longstanding unitary definition of fly sleep employed by the field, they should be doubted and debated before a consensus is reached regarding their definitions, their physiological correlates, and the best methods for their measurement. However, our results clearly establish the existence of distinct sleep states in the fly and reveal the necessity of focusing on relatively long durations of inactivity as the most sensitive means of discovering the cellular and molecular mechanisms of sleep homeostasis in *Drosophila*.

## Methods

### Flies and behavioral recording

Wildtype (*Canton-S*; BDSC stock number: 64349), *shaker[minisleep]* (*sh[mns]*; BDSC stock number: 24149), and a loss-of-function clock mutant (*per^01^*; BDSC stock number: 80928) flies were used in the experiments reported in this manuscript. Flies were maintained in bottles with Corn Syrup/Soy media (W1; Archon Scientific, Durham, NC) under 25 °C. Three to five-day-old males were collected from the bottles under CO_2_ anesthesia and loaded into glass locomotor activity tubes with sucrose-agar media. About 32 flies each were set up in *Drosophila* Activity Monitors (DAMs; Trikinetics Inc., Waltham, MA, USA) and recorded under light/dark (LD) cycles with 12-h of white light and 12-h of darkness in Tritech incubators at a constant 25 °C (Tritech Research, Los Angeles, CA, USA). During the day, the intensity of light was ∼500 lux. Flies were recorded under LD cycles for eight days, then transferred to constant darkness (DD) and temperature (25 °C) for nine days. All behavioral data were collected in 1-minute intervals/bins. Data for the three runs in Fig. 1 under LD and DD are from three different independent runs where flies were transferred to DD after exposure to LD12:12.

Also, data collected from *Canton-S* flies whose activity rhythms were recorded under UV- and Blue-blocked ramped light cycles and reported previously^31^ were used to analyze the probability of initiating sleep bouts. ∼32 flies were recorded under 25 °C and the light cycled from 10μ*W/cm*^2^ to 40-50μ*W/cm*^2^ (measured at 600nm). See Abhilash and Shafer, 2023^31^ for more details on the experimental conditions.

Data collected from *Canton-S* flies subjected to 24-h sleep deprivation using the vortexer method and a deprivation trigger frequency of 220-sec on homeostatic rebound, as previously reported^21^, were re-analyzed here to examine the extent of sleep stage-specific rebound. These flies were also recorded under LD12:12 conditions at 25 °C. See Chowdhury, Abhilash et al., 2023^21^ for more details on the experimental design. ∼32 flies were recorded per treatment. For sleep deprivation experiments, ∼32 two-to-five-day-old *sh[mns]* flies were recorded as described for the *Canton-S* flies above.

### Bout-length-based categorization of sleep states and computation of sleep bout duration

Based on the meta-analysis of studies suggesting sleep stages in flies, we chose three provisional cutoff points to segregate fly sleep (see Results). Any bout of inactivity (i.e., epoch with zero beam crossings) that was equal to or longer than 5 minutes, but less than 30 minutes, was designated “short” sleep. Any bout of inactivity that was equal to or greater than 30 minutes, but less than 60 minutes, was designated “intermediate” sleep. Finally, any bout of inactivity that was equal to or longer than 60 minutes was categorized as “long” sleep. Standard sleep was defined as any bout of inactivity that was equal to or longer than 5 minutes. For the visualization of waveform shape in Fig. 1, these sleep states were normalized to maxima. For each sleep state and each experiment, the average trace across replicate flies was normalized to the maxima, such that all the waveforms range from 0 to 1. The normalized waveforms were then averaged across replicate experiments and errors computed. This facilitates comparisons across states, experimental conditions, and studies. The waveforms of Hidden Markov Model-derived (HMM) sleep states were extracted using the online application, WebPlotDigitizer (https://wpd.starrydata2.org). Owing to data being non-normal we chose to use Kendall’s τ as the statistic to estimate correlations^56^ between our inactivity-based sleep states and the HMM-derived sleep states.

### Analysis of homeostatic sleep rebound

A schematic of the logic of analyzing sleep rebound has been shown in Fig. 4. Once “undisturbed sleep increase” is measured for each fly, we average these values. Then, we measure the “deprived sleep increase” for each fly. The difference between each fly’s “deprived sleep increase” and the averaged “undisturbed sleep increase” yields the individual experimental fly’s measure of rebound. These were calculated for standard sleep and for each of the three stages of sleep identified in this study. Statistical comparisons were made using two-tailed, one-sample *t*-tests against a known mean of 0, under the null hypothesis of no rebound in response to sleep deprivation. All analyses were carried out using custom R scripts.

Experiments with *shaker[minisleep]* (*sh[mns])* mutant flies were carried out in five independent runs. Data from all experiments were pooled for statistical testing. For each sleep state, a two-sample, two-tailed *t*-test was carried out to assess significance. Owing to there being four comparisons for the same data set, i.e., one each for standard, short, intermediate, and long sleep, we inferred statistical significance at α = 0.0125, instead of the standard Type-I error rate of 5%.

### Circular statistics of sleep states

Rose plots are circular or polar analogs of activity/sleep profiles plotted on a Cartesian (*x-y*) plane. We calculated sleep time series, binned at 30-minute intervals, under DD and UV- and Blue-blocked light cycles. A 30-minute interval was chosen as a reasonable compromise between visualizing temporal patterns of activity and sleep without losing nuance to smoothing of the data due to larger interval sizes. These were then averaged over days and across flies. The resulting average activity or sleep profiles were plotted as rose plots. Individual fly’s average activity/sleep profiles were used and transformed to polar coordinates to compute properties of the center of mass^57^. Points on a polar coordinate plane are described by *r* and θ; *r* is the distance of the point from the center of the circle and is referred to as the concentration parameter and θ is the mean angle or phase of the center of mass. These metrics have been used previously to characterize *Drosophila* rhythms^58,59^.

For each fly’s time-series of each sleep state, we computed angular deviation (s), as described previously^57^. Due to the lack of objective measurements of the onset and offset of circadian gates of sleep states, we used angular deviation as a proxy for gate-width. Angular deviation is akin to standard deviation, but in polar coordinates. It measures the angular dispersion of activity and sleep states around their mean phase. We used the onset and offset of this angular deviation as proxies of the onset and offset of behavioral sleep states. Note that in case of clearly bimodal rhythms, as determined by visual inspection of the rose plots, an angle doubling transformation was carried out before carrying out all computations^57^. Four sleep/wake states, each with an onset and offset, would result in 28 statistical comparisons (see Suppl. Tables 17-18). However, most of these comparisons are not relevant to determine the temporal architecture of the sleep states and therefore, we carried out five specific planned comparisons^60^ (see Results) and adjusted for the *p*-values by using a Benjamini-Hochberg correction, to maintain a net family-wise error rate of 5%.

### Probability of initiation of sleep bouts

For each fly and sleep state, a 1-minute timeseries was generated. From this timeseries, a list of all the standard, short, intermediate, or long sleep bouts that were initiated at each minute was curated, and the CT/ZT at which sleep bouts were initiated was noted. CT (Circadian Time) is a timescale to measure internal time. According to convention, in the fly, CT00 is defined as the time at which lights would have turned ON if the fly had continued to remain under LD cycles from which it was transferred to DD. ZT (Zeitgeber Time) is a timescale that refers to time relative to the zeitgeber (German for *time-giver*). By convention, ZT00 is defined as the time of lights-on or minimum light intensity, in the case of ramped light cycles. Once this data set was generated, we counted the number of sleep bouts that were initiated in every 1-h interval starting at CT/ZT01 and proceeding until CT/ZT24. We chose 1-h intervals to ensure sufficient sampling of bouts of individual sleep states in each time window. Then, the total number of bouts initiated in each 1-h window was divided by the total number of sleep bouts of the respective state that the fly showed. This metric serves as a measure of the frequency or probability of initiation of standard/long sleep in a 1-h window across the entire length of a 24-h cycle. For each sleep definition, i.e., standard, short, intermediate, and long sleep, a one-way repeated measures ANOVA was carried with time-point as the repeated measures variable and probability of initiation of sleep bout as the dependent variable. Post-hoc analyses were carried out using Tukey’s HSD test. All the analyses were carried out using custom R scripts. Each sleep definition under each environmental condition was analyzed using a separate ANOVA.

### Continuous Wavelet Transforms

Sleep timeseries for all three states of sleep were binned in 5-min intervals and subjected to Continuous Wavelet Transforms (CWT). A narrower temporal window was chosen in these analyses to ensure sufficiently high temporal resolution to subject our data to time series analyses. This was important, especially in the context of examining temporal variations in the strength of ultradian rhythms. Owing to biological timeseries being highly non-stationary, it was necessary to use methods such as the CWT to detect rhythmicity, as previously described^22,42–44,61^, to examine how multiple periodicities occur and change across and within circadian cycles. CWT allows for sensitively detecting rhythms across various periodicities and captures how these changes occur across timeseries^42–44,61^. The overall method of the wavelet method is like the one recently described^22^. Briefly, we used the Morlet mother wavelet to resolve our timeseries in the time-frequency domain. First, we resolved each fly’s timeseries in the time-frequency domain and then normalized it to the average amplitude of the surface. Next, we averaged the normalized surfaces across flies, which was then plotted to visualize how amplitude in different period bands vary over time (Fig. 5A). Time-averaged period versus amplitude plots were generated to capture the overall contribution of specific period bands to the overall raw timeseries of the various sleep states. A 95% Confidence Interval for the time-averaged amplitude of different period components was generated by using the bootstrap method, using 1000 replications.

To estimate the variation in amplitude/power/strength of ultradian rhythms in sleep, we considered period bands of 1 to 4 hours for short and intermediate sleep and 2 to 6 hours for long sleep. These period windows were chosen based on the signal in the scalograms for each of these sleep states (Fig. 5A). Further, owing to our definition of long sleep being 60 minutes or longer, 1-h ultradian rhythms are impossible to occur. These amplitude time-courses were then subjected to chi-squared periodogram analyses, to test for rhythmicity in ultradian rhythm amplitudes in each of the sleep states.

### Introducing phaseR

*phaseR* is an RStudio and Shiny-based application that we have developed to facilitate all the analyses we report in this manuscript. It enables the analysis of user-defined sleep stages, and the application of wavelet transforms to activity and sleep time series. The application can be downloaded from its GitHub repository (*Abhilash and Shafer, manuscript in preparation*; https://github.com/abhilashlakshman/phaseR). Most functions used in *phaseR* are part of an R package, called *phase*^62^. Functions that are not part of *phase*, have been implemented in the server environment of the Shiny application. Please see the user manual for a detailed description of all the functionalities and instructions on how to use them.

## Acknowledgements

We thank Hannah Pettibone, Matthew Ciolkowski, Budhaditya Chowdhury, William Hunter Bauserman, and Jacqueline Jou for assistance with experiments and logistics. We are grateful to Benjamin Smarr for several discussions on wavelet analyses that have been extensively used in this study. We thank María de la Paz Fernández and two anonymous reviewers for useful feedback on a previous version of this manuscript. This work was supported by grants from the National Institute of Neurological Disorders and Stroke (R01NS077933 and R21NS131939).

