## Supplemental Tables and Figures for "Recognition of distinct sleep states in *Drosophila* uncovers previously obscured homeostatic and circadian control of sleep"

*Running title: Drosophila sleep states*

**Suppl. Table 1:** Results of correlations between our provisional sleep states and sleep states inferred through Hidden Markov Models. Kendall's tau correlations were carried out. Reported here are test statistics and *p*-values for each comparison. Also see Fig. 1.

| | **Behavioral states under LD** | **Behavioral states under DD** |
|---|---|---|
| Short Sleep *vs* Early Wake | $z = 2.87; p = 0.0041$ | $z = 3.16; p = 0.0016$ |
| Inter. Sleep *vs* Light Sleep | $z = 2.73; p = 0.0063$ | $z = 2.27; p = 0.0234$ |
| Long Sleep *vs* Deep Sleep | $z = 2.92; p = 0.0035$ | $z = 5.07; p < 0.0001$ |

**Suppl. Table 2:** Results of the repeated measures ANOVA to test the effect of time-point on the probability of initiating *standard* sleep under LD cycles, in wildtype flies, under 25 °C. Significant effects are italicized and marked in red. A statistically significant effect of time-point is taken to indicate the presence of a rhythm. Also see Fig. 2.

| | *df Effect* | *MS Effect* | *df Error* | *MS Error* | *F* | *p* |
|---|---|---|---|---|---|---|
| ***Time-point*** | *23* | *0.00663* | *713* | *0.0006* | *11.14* | *<2e-16* |
| ***Channel*** | 31 | 1.05e-19 | | | | |

**Suppl. Table 3:** Results of the repeated measures ANOVA to test the effect of time-point on the probability of initiating *short* sleep under LD cycles, in wildtype flies, under 25 °C. Significant effects are italicized and marked in red. A statistically significant effect of time-point is taken to indicate the presence of a rhythm. Also see Fig. 2.

| | *df Effect* | *MS Effect* | *df Error* | *MS Error* | *F* | *p* |
|---|---|---|---|---|---|---|
| ***Time-point*** | *23* | *0.011* | *713* | *0.00102* | *10.75* | *<2e-16* |
| ***Channel*** | 31 | 1.40e-19 | | | | |

**Suppl. Table 4:** Results of the repeated measures ANOVA to test the effect of time-point on the probability of initiating *intermediate* sleep under LD cycles, in wildtype flies, under 25 °C. Significant effects are italicized and marked in red. A statistically significant effect of time-point is taken to indicate the presence of a rhythm. Also see Fig. 2.

| | *df Effect* | *MS Effect* | *df Error* | *MS Error* | *F* | *p* |
|---|---|---|---|---|---|---|
| ***Time-point*** | *23* | *0.01277* | *713* | *0.00189* | *6.74* | *<2e-16* |
| ***Channel*** | 31 | 2.14e-19 | | | | |

**Suppl. Table 5:** Results of the repeated measures ANOVA to test the effect of time-point on the probability of initiating *long* sleep under LD cycles, in wildtype flies, under 25 °C. Significant effects are italicized and marked in red. A statistically significant effect of time-point is taken to indicate the presence of a rhythm. Also see Fig. 2.

| | *df effect* | *MS effect* | *df Error* | *MS Error* | *F* | *p* |
|---|---|---|---|---|---|---|
| ***Time-point*** | *23* | *0.0248* | *713* | *0.00189* | *13.16* | *<2e-16* |
| ***Channel*** | 31 | 1.05e-19 | | | | |

**Suppl. Table 6:** Values of probabilities (Avg. Probs.) of initiating standard sleep and each of the sleep stages at different times of the day under LD cycles, compiled from the raw data on which the ANOVAs above were carried out. Post hoc multiple comparisons were carried out using the Tukey's Honestly Significant Difference (HSD) test. Time-points that share letters in the "Groups" column were not significantly different from each other. Also see Fig. 2.

| | **Std. sleep** | | **Short sleep** | | **Inter. sleep** | | **Long sleep** | |
|---|---|---|---|---|---|---|---|---|
| *ZT* | *Avg. Probs.* | *Groups* | *Avg. Probs.* | *Groups* | *Avg. Probs.* | *Groups* | *Avg. Probs.* | *Groups* |
| *1* | 0.0469 | bcdef | 0.0543 | bc | 0.0170 | ef | 0.0412 | bc |
| *2* | 0.0796 | a | 0.0898 | a | 0.0365 | bcdef | 0.0636 | b |
| *3* | 0.0522 | bcd | 0.0488 | bcd | 0.0619 | abc | 0.0599 | b |
| *4* | 0.0359 | defg | 0.0311 | bcde | 0.0367 | bcdef | 0.0553 | b |
| *5* | 0.0307 | defg | 0.0238 | de | 0.0380 | bcdef | 0.0596 | b |
| *6* | 0.0245 | gh | 0.0211 | de | 0.0198 | def | 0.0446 | bc |
| *7* | 0.0272 | fgh | 0.0254 | cde | 0.0294 | bcdef | 0.0327 | bcd |
| *8* | 0.0283 | efgh | 0.0286 | bcde | 0.0261 | cdef | 0.0366 | bcd |
| *9* | 0.0340 | defg | 0.0375 | bcde | 0.0581 | abcd | 0.0074 | cd |
| *10* | 0.0629 | ab | 0.0906 | a | 0.0198 | def | 0.0009 | d |
| *11* | 0.0329 | defg | 0.0494 | bcd | 0.0022 | f | 0.0000 | d |
| *12* | 0.0075 | h | 0.0106 | e | 0.0026 | f | 0.0000 | d |
| *13* | 0.0626 | abc | 0.0475 | bcd | 0.0372 | bcdef | 0.1282 | a |
| *14* | 0.0406 | cdefg | 0.0365 | bcde | 0.0542 | abcde | 0.0410 | bc |
| *15* | 0.0381 | defg | 0.0343 | bcde | 0.0428 | abcde | 0.0427 | bc |
| *16* | 0.0415 | bcdefg | 0.0371 | bcde | 0.0478 | abcde | 0.0564 | b |
| *17* | 0.0455 | bcdefg | 0.0410 | bcd | 0.0579 | abcd | 0.0452 | bc |
| *18* | 0.0425 | bcdefg | 0.0378 | bcde | 0.0510 | abcde | 0.0533 | b |
| *19* | 0.0393 | defg | 0.0301 | bcde | 0.0562 | abcde | 0.0589 | b |
| *20* | 0.0407 | bcdefg | 0.0343 | bcde | 0.0486 | abcde | 0.0557 | b |
| *21* | 0.0434 | bcdefg | 0.0394 | bcde | 0.0614 | abc | 0.0442 | bc |
| *22* | 0.0479 | bcdef | 0.0442 | bcd | 0.0802 | a | 0.0371 | bcd |
| *23* | 0.0499 | bcde | 0.0488 | bcd | 0.0675 | ab | 0.0345 | bcd |
| *24* | 0.0455 | bcdefg | 0.0578 | b | 0.0472 | abcde | 0.0010 | d |

**Suppl. Table 7:** Results of the repeated measures ANOVA to test the effect of time-point on the probability of initiating *standard* sleep under constant darkness, in wildtype flies, under 25 °C. Significant effects are italicized and marked in red. A statistically significant effect of time-point is taken to indicate the presence of a rhythm. Also see Fig. 2.

| | *df Effect* | *MS Effect* | *df Error* | *MS Error* | *F* | *p* |
|---|---|---|---|---|---|---|
| ***Time-point*** | *23* | *0.0035* | *690* | *0.00023* | *15.06* | *<2e-16* |
| ***Channel*** | 30 | 1.39e-19 | | | | |

**Suppl. Table 8:** Results of the repeated measures ANOVA to test the effect of time-point on the probability of initiating *short* sleep under constant darkness, in wildtype flies, under 25 °C. Significant effects are italicized and marked in red. A statistically significant effect of time-point is taken to indicate the presence of a rhythm. Also see Fig. 2.

| | *df Effect* | *MS Effect* | *df Error* | *MS Error* | *F* | *p* |
|---|---|---|---|---|---|---|
| ***Time-point*** | *23* | *0.00844* | *690* | *0.00047* | *18.05* | *<2e-16* |
| ***Channel*** | 30 | 2.38e-19 | | | | |

**Suppl. Table 9:** Results of the repeated measures ANOVA to test the effect of time-point on the probability of initiating *intermediate* sleep under constant darkness, in wildtype flies, under 25 °C. Significant effects are italicized and marked in red. A statistically significant effect of time-point is taken to indicate the presence of a rhythm. Also see Fig. 2.

| | *df Effect* | *MS Effect* | *df Error* | *MS Error* | *F* | *p* |
|---|---|---|---|---|---|---|
| ***Time-point*** | *23* | *0.0056* | *690* | *0.00127* | *4.416* | *6.38e-11* |
| ***Channel*** | 30 | 2.86e-19 | | | | |

**Suppl. Table 10:** Results of the repeated measures ANOVA to test the effect of time-point on the probability of initiating *long* sleep under constant darkness, in wildtype flies, under 25 °C. Significant effects are italicized and marked in red. A statistically significant effect of time-point is taken to indicate the presence of a rhythm. Also see Fig. 2.

| | *df Effect* | *MS Effect* | *df Error* | *MS Error* | *F* | *p* |
|---|---|---|---|---|---|---|
| ***Time-point*** | *23* | *0.01372* | *667* | *0.00204* | *6.732* | *<2e-16* |
| ***Channel*** | 29 | 1.91e-19 | | | | |

**Suppl. Table 11:** Values of probabilities (Avg. Probs.) of initiating standard sleep and each of the sleep stages at different times of the day under constant darkness, compiled from the raw data on which the ANOVAs above were carried out. Post hoc multiple comparisons were carried out using the Tukey's Honestly Significant Difference (HSD) test. Time-points that share letters in the "Groups" column were not significantly different from each other. Also see Fig. 2.

| | **Std. sleep** | | **Short sleep** | | **Inter. sleep** | | **Long sleep** | |
|---|---|---|---|---|---|---|---|---|
| *CT* | *Avg. Probs.* | *Groups* | *Avg. Probs.* | *Groups* | *Avg. Probs.* | *Groups* | *Avg. Probs.* | *Groups* |
| *1* | 0.0486 | abcd | 0.0444 | cdef | 0.0603 | a | 0.0368 | bcdef |
| *2* | 0.0477 | abcd | 0.0453 | bcde | 0.0573 | ab | 0.0313 | cdef |
| *3* | 0.0507 | abc | 0.0492 | abcd | 0.0596 | a | 0.0451 | abcdef |
| *4* | 0.0494 | abcd | 0.0459 | bcd | 0.0588 | a | 0.0411 | bcdef |
| *5* | 0.0473 | abcde | 0.0454 | bcde | 0.0571 | ab | 0.0284 | def |
| *6* | 0.0532 | abc | 0.0568 | abc | 0.0604 | a | 0.0217 | ef |
| *7* | 0.0576 | a | 0.0647 | ab | 0.0461 | abcd | 0.0231 | ef |
| *8* | 0.0561 | ab | 0.0663 | a | 0.0427 | abcd | 0.0118 | f |
| *9* | 0.0547 | ab | 0.0680 | a | 0.0249 | bcd | 0.0126 | ef |
| *10* | 0.0513 | abc | 0.0669 | a | 0.0229 | cd | 0.0097 | f |
| *11* | 0.0473 | abcde | 0.0586 | abc | 0.0176 | d | 0.0214 | ef |
| *12* | 0.0422 | bcdefg | 0.0488 | abcd | 0.0191 | cd | 0.0250 | ef |
| *13* | 0.0435 | abcdefg | 0.0439 | cdef | 0.0243 | bcd | 0.0494 | abcdef |
| *14* | 0.0336 | efghi | 0.0296 | defgh | 0.0400 | abcd | 0.0489 | abcdef |
| *15* | 0.0308 | fghi | 0.0258 | efgh | 0.0350 | abcd | 0.0707 | abcd |
| *16* | 0.0263 | hi | 0.0172 | h | 0.0371 | abcd | 0.0710 | abcd |
| *17* | 0.0247 | i | 0.0180 | h | 0.0384 | abcd | 0.0545 | abcde |
| *18* | 0.0258 | i | 0.0194 | h | 0.0319 | abcd | 0.0844 | a |
| *19* | 0.0277 | hi | 0.0209 | gh | 0.0431 | abcd | 0.0714 | abc |
| *20* | 0.0298 | ghi | 0.0250 | fgh | 0.0396 | abcd | 0.0763 | ab |
| *21* | 0.0316 | fghi | 0.0255 | efgh | 0.0453 | abcd | 0.0472 | abcdef |
| *22* | 0.0357 | defghi | 0.0321 | defgh | 0.0471 | abcd | 0.0468 | abcdef |
| *23* | 0.0400 | cdefgh | 0.0405 | cdefg | 0.0407 | abcd | 0.0341 | bcdef |
| *24* | 0.0440 | abcdef | 0.0417 | cdef | 0.0508 | abc | 0.0372 | bcdef |

**Suppl. Table 12:** Results of the repeated measures ANOVA to test the effect of time-point on the probability of initiating *standard* sleep under UV- and Blue-blocked ramped light cycles, in wildtype flies, under 25 °C. Significant effects are italicized and marked in red. A statistically significant effect of time-point is taken to indicate the presence of a rhythm. Also see Fig. 2.

| | *df Effect* | *MS Effect* | *df Error* | *MS Error* | *F* | *p* |
|---|---|---|---|---|---|---|
| ***Time-point*** | *23* | *0.00456* | *621* | *0.00059* | *7.757* | *<2e-16* |
| ***Channel*** | 27 | 2.22e-19 | | | | |

**Suppl. Table 13:** Results of the repeated measures ANOVA to test the effect of time-point on the probability of initiating *short* sleep under UV- and Blue-blocked ramped light cycles, in wildtype flies, under 25 °C. Significant effects are italicized and marked in red. A statistically significant effect of time-point is taken to indicate the presence of a rhythm. Also see Fig. 2.

| | *df Effect* | *MS Effect* | *df Error* | *MS Error* | *F* | *p* |
|---|---|---|---|---|---|---|
| ***Time-point*** | *23* | *0.00525* | *621* | *0.00099* | *5.287* | *7.73e-14* |
| ***Channel*** | 27 | 1.03e-19 | | | | |

**Suppl. Table 14:** Results of the repeated measures ANOVA to test the effect of time-point on the probability of initiating *intermediate* sleep under UV- and Blue-blocked ramped light cycles, in wildtype flies, under 25 °C. Significant effects are italicized and marked in red. A statistically significant effect of time-point is taken to indicate the presence of a rhythm. Also see Fig. 2.

| | *df Effect* | *MS Effect* | *df Error* | *MS Error* | *F* | *p* |
|---|---|---|---|---|---|---|
| ***Time-point*** | *23* | *0.01065* | *621* | *0.00402* | *2.65* | *5.14e-05* |
| ***Channel*** | 27 | 1.23e-19 | | | | |

**Suppl. Table 15:** Results of the repeated measures ANOVA to test the effect of time-point on the probability of initiating *long* sleep under UV- and Blue-blocked ramped light cycles, in wildtype flies, under 25 °C. Significant effects are italicized and marked in red. A statistically significant effect of time-point is taken to indicate the presence of a rhythm. Also see Fig. 2.

| | *df Effect* | *MS Effect* | *df Error* | *MS Error* | *F* | *p* |
|---|---|---|---|---|---|---|
| ***Time-point*** | *23* | *0.03509* | *621* | *0.00562* | *6.246* | *<2e-16* |
| ***Channel*** | 27 | 6.59e-20 | | | | |

**Suppl. Table 16:** Values of probabilities (Avg. Probs.) of initiating standard sleep and each of the sleep stages at different times of the day under UV- and Blue-blocked ramped light cycles, compiled from the raw data on which the ANOVAs above were carried out. Post hoc multiple comparisons were carried out using the Tukey's Honestly Significant Difference (HSD) test. Time-points that share letters in the "Groups" column were not significantly different from each other. Also see Fig. 2.

| | **Std. sleep** | | **Short sleep** | | **Inter. sleep** | | **Long sleep** | |
|---|---|---|---|---|---|---|---|---|
| *ZT* | *Avg. Probs.* | *Groups* | *Avg. Probs.* | *Groups* | *Avg. Probs.* | *Groups* | *Avg. Probs.* | *Groups* |
| *1* | 0.0332 | bcdef | 0.0223 | defg | 0.0518 | ab | 0.1073 | ab |
| *2* | 0.0270 | cdef | 0.0255 | cdefg | 0.0217 | ab | 0.0414 | bcde |
| *3* | 0.0349 | bcdef | 0.0305 | bcdefg | 0.0560 | ab | 0.0317 | cde |
| *4* | 0.0398 | bcde | 0.0397 | abcdefg | 0.0431 | ab | 0.0156 | de |
| *5* | 0.0473 | abc | 0.0487 | abcdefg | 0.0486 | ab | 0.0455 | bcde |
| *6* | 0.0498 | abc | 0.0508 | abcde | 0.0624 | ab | 0.0197 | de |
| *7* | 0.0657 | a | 0.0656 | a | 0.0651 | ab | 0.0487 | bcde |
| *8* | 0.0549 | ab | 0.0523 | abcd | 0.0575 | ab | 0.0680 | abcde |
| *9* | 0.0492 | abc | 0.0467 | abcdefg | 0.0624 | ab | 0.0373 | bcde |
| *10* | 0.0524 | ab | 0.0490 | abcdef | 0.0574 | ab | 0.0453 | bcde |
| *11* | 0.0472 | abc | 0.0443 | abcdefg | 0.0541 | ab | 0.0515 | bcde |
| *12* | 0.0506 | abc | 0.0464 | abcdefg | 0.0691 | a | 0.0456 | bcde |
| *13* | 0.0470 | abc | 0.0460 | abcdefg | 0.0561 | ab | 0.0364 | bcde |
| *14* | 0.0483 | abc | 0.0506 | abcde | 0.0523 | ab | 0.0271 | de |
| *15* | 0.0552 | ab | 0.0613 | ab | 0.0504 | ab | 0.0268 | de |
| *16* | 0.0487 | abc | 0.0559 | abc | 0.0321 | ab | 0.0113 | de |
| *17* | 0.0433 | abcd | 0.0528 | abcd | 0.0256 | ab | 0.0030 | de |
| *18* | 0.0369 | bcdef | 0.0485 | abcdefg | 0.0212 | ab | 0.0000 | e |
| *19* | 0.0208 | def | 0.0251 | cdefg | 0.0060 | b | 0.0090 | de |
| *20* | 0.0156 | f | 0.0181 | g | 0.0126 | ab | 0.0030 | de |
| *21* | 0.0166 | ef | 0.0198 | fg | 0.0062 | b | 0.0095 | de |
| *22* | 0.0463 | abc | 0.0439 | abcdefg | 0.0186 | ab | 0.0758 | abcd |
| *23* | 0.0410 | bcd | 0.0347 | bcdefg | 0.0341 | ab | 0.1379 | a |
| *24* | 0.0282 | cdef | 0.0215 | efg | 0.0356 | ab | 0.1027 | abc |

**Suppl. Table 17:** Statistical comparisons of phases of different sleep/wake states under constant darkness. Planned one-tailed paired Wilcoxon's tests were carried out (see text for details). Reported are *p*-values adjusted for a family-wise error rate of 5% using the Benjamini-Hochberg correction, and n.t. indicate pairs that were not tested. Green highlighted cells are the planned comparisons that were carried out. Pairs with significant differences are italicized and in red. Also see Fig. 3.

| | Act. onset | Act. offset | Long onset | Long offset | Inter. onset | Inter. offset | Short onset | Short offset |
|---|---|---|---|---|---|---|---|---|
| Act. onset | | | | | | | | |
| Act. offset | n.t. | | | | | | | |
| Long onset | n.t. | *<0.01* | | | | | | |
| Long offset | n.t. | n.t. | n.t. | | | | | |
| Inter. onset | n.t. | n.t. | n.t. | *<0.001* | | | | |
| Inter. offset | *<0.01* | n.t. | n.t. | n.t. | n.t. | | | |
| Short onset | n.t. | n.t. | n.t. | *<0.001* | n.t. | n.t. | | |
| Short offset | 0.95 | n.t. | n.t. | n.t. | n.t. | n.t. | n.t. | |

**Suppl. Table 18:** Statistical comparisons of phases of different sleep/wake states under UV- and Blue-blocked ramped light cycles. Planned one-tailed paired Wilcoxon's tests were carried out (see text for details). Reported are *p*-values adjusted for a family-wise error rate of 5% using the Benjamini-Hochberg correction, and n.t. indicate pairs that were not tested. Green highlighted cells are the planned comparisons that were carried out. Pairs with significant differences are italicized and in red. Also see Fig. 3.

| | Act. onset | Act. offset | Long onset | Long offset | Inter. onset | Inter. offset | Short onset | Short offset |
|---|---|---|---|---|---|---|---|---|
| Act. onset | | | | | | | | |
| Act. offset | n.t. | | | | | | | |
| Long onset | n.t. | *<0.001* | | | | | | |
| Long offset | n.t. | n.t. | n.t. | | | | | |
| Inter. onset | n.t. | n.t. | n.t. | 0.03 | | | | |
| Inter. offset | *<0.001* | n.t. | n.t. | n.t. | n.t. | | | |
| Short onset | n.t. | n.t. | n.t. | *<0.01* | n.t. | n.t. | | |
| Short offset | *<0.001* | n.t. | n.t. | n.t. | n.t. | n.t. | n.t. | |

**Suppl. Table 19:** Results of chi-square periodogram analyses to test rhythmicity in the power of ultradian rhythms in the three different sleep states. Also see Fig. 6.

| | ***wildtype*** | | | ***loss-of-function clock mutant*** | | |
|---|---|---|---|---|---|---|
| | *%rhythmic* | *Period (h)* | *Power* | *%rhythmic* | *Period (h)* | *Power* |
| *Short sleep* | *96.77* | 23.44 | 389.24 | 32.14 | 23.85 | 144.66 |
| *Inter. sleep* | *100.00* | 23.41 | 383.32 | 32.14 | 24.85 | 62.74 |
| *Long sleep* | *93.33* | 23.83 | 424.73 | 24.33 | 24.33 | 110.91 |

**Suppl. Fig. 1:** Averaged timeseries of sleep states before, during and after mechanical sleep deprivation. Also see Fig. 4.

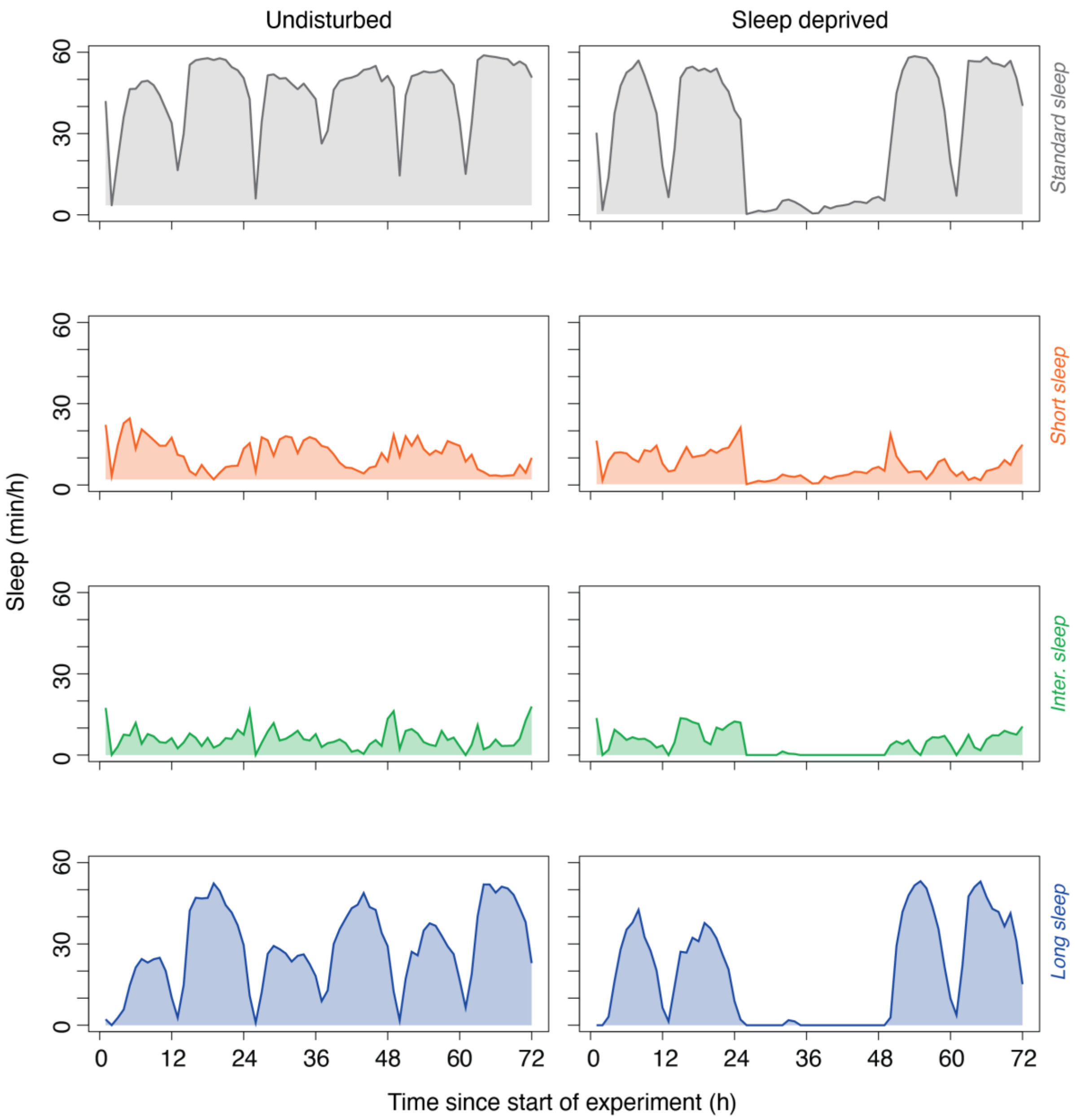

**Suppl. Fig. 2:** Averaged timeseries of sleep states before, during and after mechanical sleep deprivation for the classic sleep homeostat mutant, *shaker[minisleep]*. Each trace is the average waveform for each replicate run. Gray shaded regions indicate the dark phase of the light/dark cycle. Also see Fig. 4.

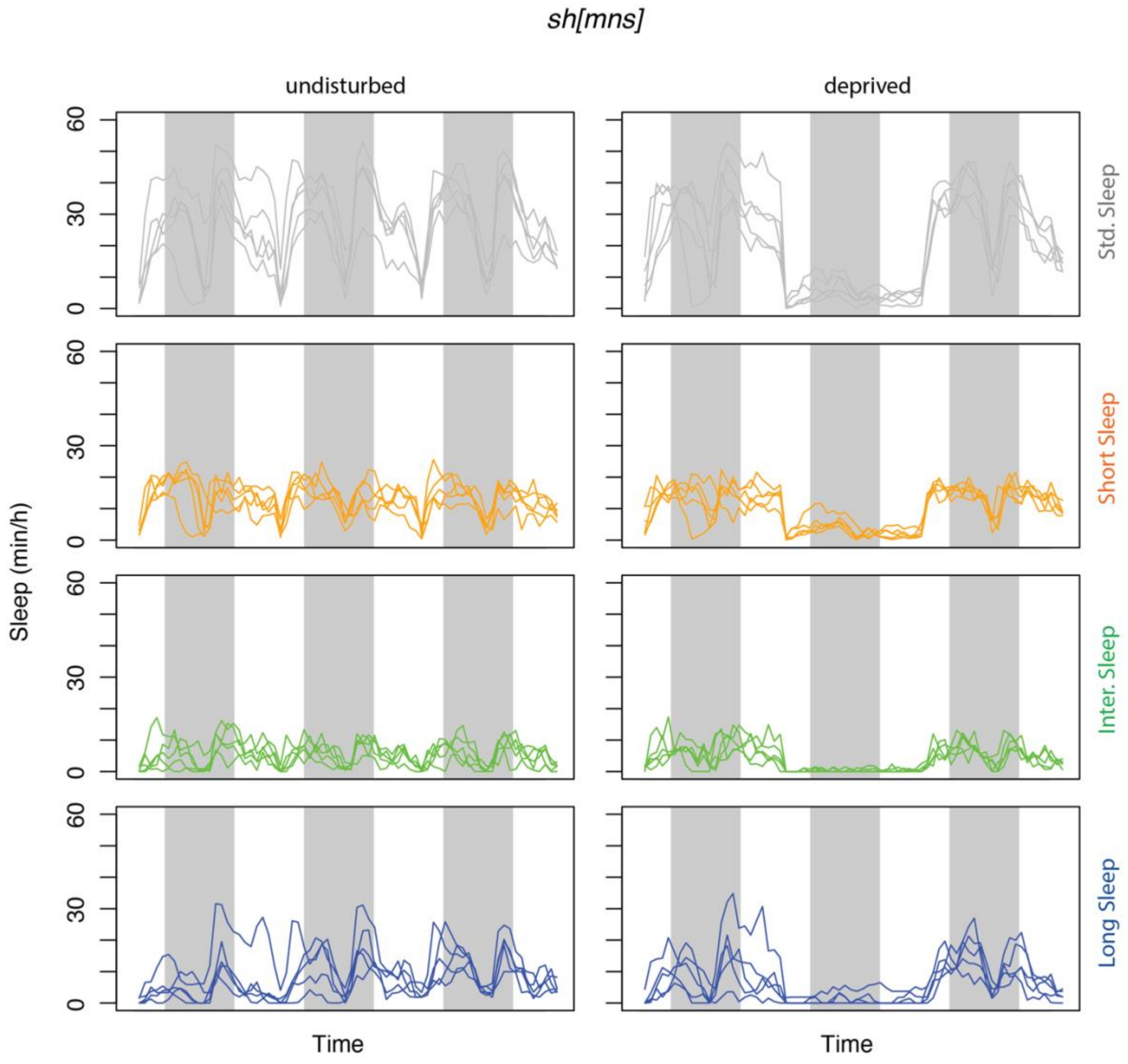

**Suppl. Fig. 3:** Schematic illustrating the distinction between measurements of sleep fragmentation/consolidation and sleep staging. In this example, the amount of long sleep is not changed despite substantial changes in average bout length, the commonly used metric to assess fragmentation/consolidation.

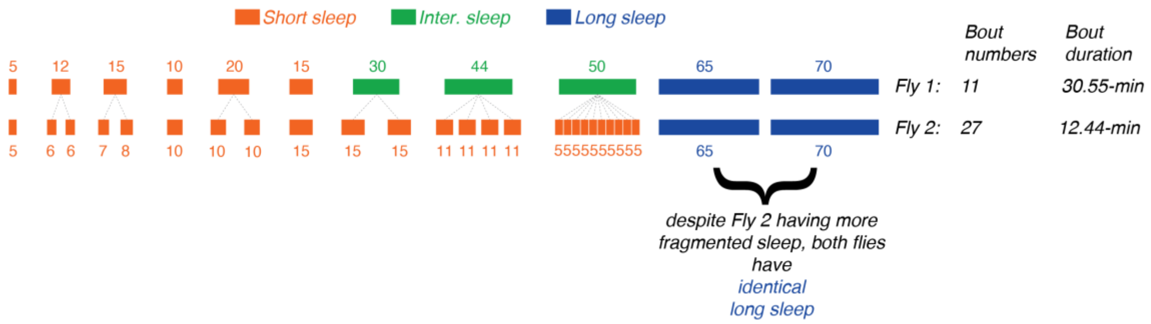

**Suppl. Fig. 4:** A single fly's representative time course to demonstrate that although each sleep state cannot co-exist in time, it is possible for their circadian gates to overlap. Light and dark gray shaded areas represent the subjective day and night phases, respectively. Also see Fig. 3.

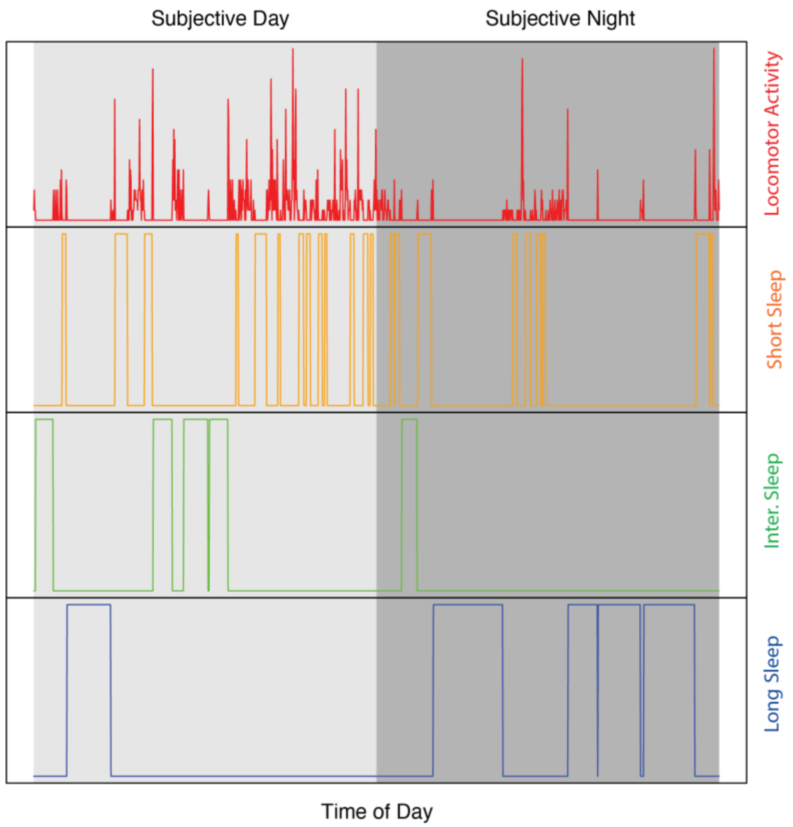
